# Structural diversity of mitochondria in the neuromuscular system across development

**DOI:** 10.1101/2024.07.19.604219

**Authors:** J. Alexander Bae, Myung-kyu Choi, Soungyub Ahn, Gwanho Ko, Daniel T. Choe, Hyunsoo Yim, Ken C. Nguyen, Jinseop S. Kim, David H. Hall, Junho Lee

**Affiliations:** Research Institute of Basic Sciences, Seoul National University, Seoul, Republic of Korea; Department of Biological Sciences, Seoul National University, Seoul, Republic of Korea; Dominick P. Purpura Department of Neuroscience, Albert Einstein College of Medicine, Bronx, NY, USA; Department of Structure and Function of Neural Networks, Korea Brain Research Institute, Daegu, Republic of Korea; Department of Biological Sciences, Sungkyunkwan University, Suwon, Republic of Korea

## Abstract

As an animal matures, its neural circuit undergoes alterations, yet the developmental changes in intracellular organelles to facilitate these changes is less understood. Using 3D electron microscopy and deep learning, we developed semi-automated methods for reconstructing mitochondria in *C*. *elegans* and collected mitochondria reconstructions from normal reproductive stages and dauer, enabling comparative study on mitochondria structure within the neuromuscular system. We found that various mitochondria structural properties in neurons correlate with synaptic connections and these properties are preserved across development in different neural circuits. To test the necessity of these universal mitochondria properties, we examined the behavior in *drp-1* mutants with impaired mitochondria fission and discovered that it caused behavioral deficits. Moreover, we observed that dauer neurons display distinctive mitochondrial features, and mitochondria in dauer muscles exhibit unique reticulum-like structure. We propose that this specialized mitochondria structure may serve as an adaptive mechanism to support stage-specific behavioral and physiological needs.

## Introduction

The neural circuit undergoes continuous changes throughout development in various brain regions, adapting for more complex computations and to perform more resilient behavior (*Neural Circuit Development and Function in the Healthy and Diseased Brain: Comprehensive Developmental Neuroscience* 2013; Tau and Peterson 2010). This functional adaptation across development is accompanied by the structural alterations, resulting in changes to both neuronal morphology and connectivity between neurons (Grueber et al. 2005; Kroon et al. 2019; Khalil, Farhat, and Dłotko 2021; Kolk and Rakic 2022; Witvliet et al. 2021; Mulcahy et al. 2022). To induce changes in neuronal morphology and connectivity, numerous intracellular organelles participate in the process, with mitochondria playing a central role through trafficking and reshaping (Harris, Jolivet, and Attwell 2012; Verstreken et al. 2005; Kimura and Murakami 2014; Sheng and Cai 2012; Courchet et al. 2013). However, less is known about how mitochondria structure evolves throughout development to support diverse neural circuits and cellular processes.

3D electron microscopy (EM) provides a distinctive advantage in studying structural properties of mitochondria due to its superior resolution. It allows us to study not only the detailed morphology of individual mitochondrion but also the overall organization of all mitochondria in the volume. Besides, it enables us to relate the mitochondria structural properties with other cellular features such as neurite morphology and synapses since it contains various structural information.

Previously, a number of studies investigated mitochondria structure using 3D EM in mammalian neurons (Kasthuri et al. 2015; Smith et al. 2016; Bloss et al. 2018; Calì et al. 2018; Turner et al. 2022). However, as mammalian neural circuits are quite large, most studies were limited to a small number of neurons or to selected regions of the brain. In invertebrates, a comprehensive study on mitochondria structure in *Drosophila* has been reported recently; however, this study did not include analysis of mitochondria in muscle cells (Rivlin et al. 2024). In order to gain deeper insights into the connection between mitochondria structure and neural circuit function, it is essential to investigate mitochondria structure within an entire system from sensory neurons to muscle cells as there could be regional differences in the neuromuscular system (White et al. 1986).

Besides, most EM studies on mitochondria are limited to single time point (Kasthuri et al. 2015; Smith et al. 2016; Bloss et al. 2018; Turner et al. 2022; Rivlin et al. 2024). When multiple time points are examined, the studies compare only a few broadly defined intervals like young adult vs. aged (Calì et al. 2018). However, these time points are rather ill-defined, making it difficult to identify stereotypic cellular or circuit functions at each interval. In contrast, *C*. *elegans* developmental stages are clearly distinguishable, and numerous studies have reported findings on stage-specific cellular physiology and behavior in an alternative developmental stage (Cassada and Russell 1975; Golden and Riddle 1984a, 1984b; Gaglia and Kenyon 2009; Hallem et al. 2011; Wolkow and Hall 2016; H. Lee et al. 2011).

Here we use 3D EM and *C*. *elegans* as a model organism to answer the above questions as it becomes feasible to conduct a comprehensive analysis of mitochondria structure within an entire system across development. We have densely reconstructed mitochondria from 3D EM images in the *C*. *elegans* body near the nerve ring for various developmental stages from L1 to adult, and dauer stage (Figure 1; Witvliet et al. 2021; Yim et al. 2024). Our EM reconstructions provide us with opportunities to study detailed mitochondria structure in both neurons and muscle cells across multiple developmental stages from the initial L1 stage after the egg hatch to adult, and also the alternative long-lived stage, the dauer. As a result, we found that there exists fundamental principles in mitochondria structure that are preserved across development. In addition, we have tested that this specific structure is necessary to operate the neural circuit resulting in the intended behavior. Lastly, we observed that mitochondria in the dauer stage exhibit distinct morphology in both neuronal and non-neuronal cells, providing insights into their role during the energy-conserving phase.

**Figure 1.**
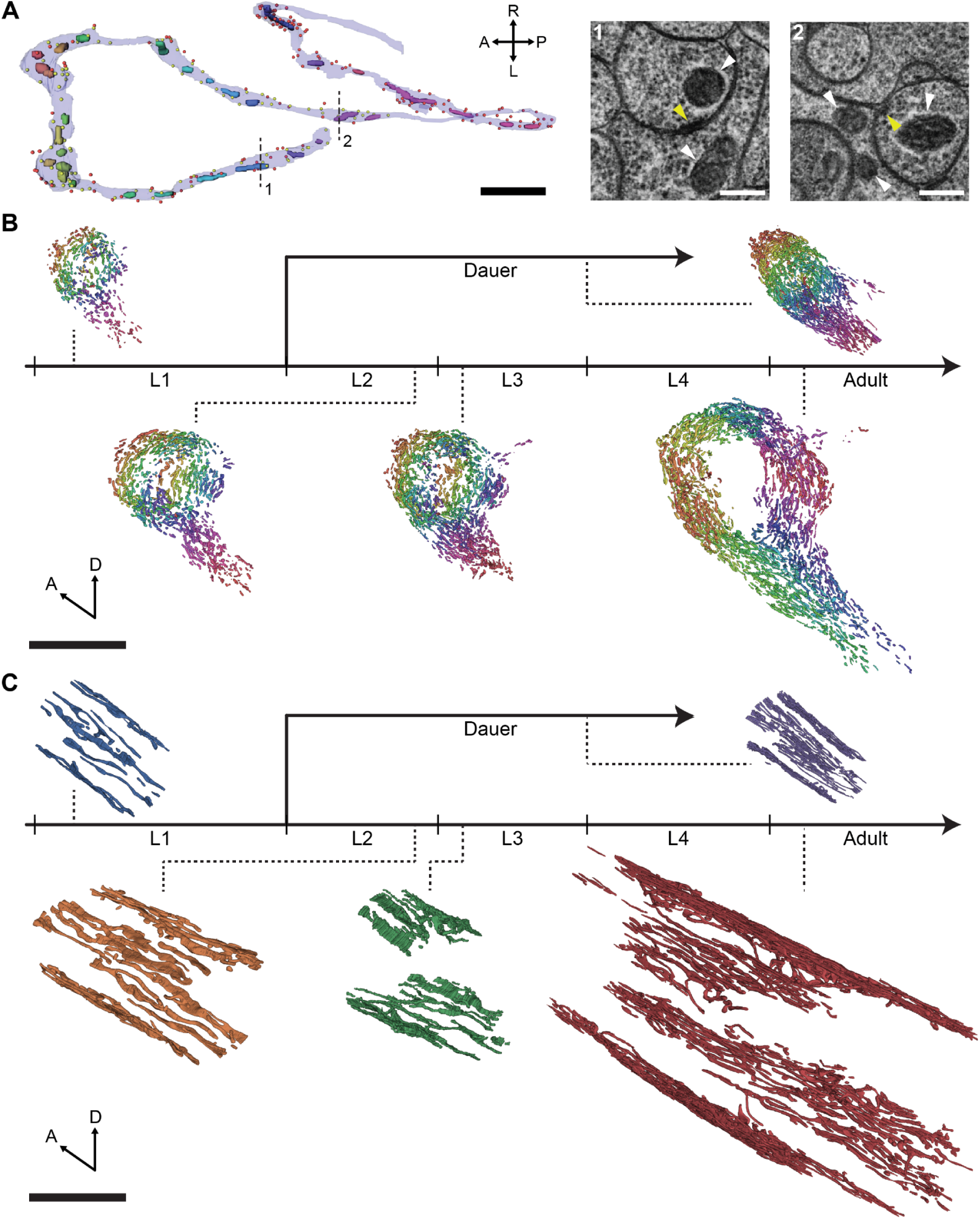
Comprehensive mitochondria reconstructions of *C*. *elegans* across development. (A) Mitochondria reconstructions in a sample neuron, RIAR, in the dauer stage with presynaptic sites (i.e., active zones; yellow circles) and postsynaptic sites (red circles) labeled (left). Parts of thin sections at the locations indicated by dashed lines (right) including mitochondria (white arrows) and active zones (yellow arrows). (B and C) Dense neuronal mitochondria (B) and body wall muscle mitochondria (C) reconstructions in different developmental stages: L1, L2, L3, adult, and dauer. (A) Scale bars: 2 µm (black), 200 nm (white). (B and C) Scale bar: 10 µm. (A and B) Different mitochondria are denoted by different colors. (C) Colors of mitochondria indicate their stage. See also Figure S1.

## Results

### Comprehensive mitochondria reconstructions across development

Mitochondria have distinctive visual features in EM (Figure 1A) so it is possible to visually identify mitochondria in EM. Consequently, we have trained a convolutional neural network (CNN) to automatically detect mitochondria in each image section (Figure S1A, Methods). Then, we have reconstructed mitochondria in 3D by combining predictions from adjacent sections (Figure S1A, Methods).

We have densely reconstructed mitochondria in different developmental stages of *C*. *elegans*: L1, L2, L3, adult, and dauer (Figures 1B and 1C). For normal reproductive stages, we have used publicly available EM image volumes which have been published recently (Witvliet et al. 2021). For dauer, we imaged the *C*. *elegans* dauer with length of ∼18 μm using serial-section EM (Yim et al. 2024). We have reconstructed all the mitochondria, both in neuronal and non-neuronal cells like body wall muscles (BWMs), included in the EM volume (Figures 1B and 1C).

In the neurites of neurons, mitochondria exist in pill-shaped pieces and they rarely have branching (Figure 1B). The total number of mitochondria in neurons had a linearly increasing trend from L1 to adulthood, approximately a five-fold increase (Figure S1B; L1: n=452, L2: n=763; L3: n=1009, Adult: n=1775). This is similar to the increase in neurite length and body length from L1 to adulthood (Witvliet et al. 2021). In the dauer stage, neurons contained about 20% more mitochondria than neurons in L3 (n=1227), which is a similar stage in terms of developmental cycle (Figure S1B).

The size of mitochondria increased from L1 to L2, then the size of mitochondria saturated to a similar level from L2 to adult (Figure S1C). The dauer neurons contained substantially smaller mitochondria while the lengths of mitochondria were comparable (Figures S1C and S1D). As more and larger mitochondria exist in adult neurons due to their larger volume, mitochondria in the nerve ring showed eight-fold increase in total volume (Figures S1E-S1G). The dauer stage showed similar value with stage L2 (Figure S1E).

In the BWMs, mitochondria exist in elongated form spanning over the muscles, which is expected as the mitochondria need to provide energy in every location along the body. As the worm develops from L1 to adult, the mitochondria form more branches leading to more complex shape with aligned strands, similar to the previous report (Figure 1C; Han et al. 2012; Schultz et al. 2017). The BWM mitochondria in dauer has more branching as in adults but are composed of sparser strands which we will elaborate in the later section (Figure 1C).

### Mitochondria structure is related to synaptic connections

Mitochondria are known to be closely associated with synaptic connections. Since we are provided with both the morphology and spatial locations of synapses and mitochondria, it is possible to investigate the relation between them. As a result, we found out-degree, the number of outgoing synapses, is correlated with the number of mitochondria (n=161, *r*=0.65, *p*<10^−19^; Pearson correlation) and the total amount of mitochondria (n=161, *r*=0.62, *p*<10^−17^; Pearson correlation) in a neuron in all developmental stages (Figures 2A and S2A). Moreover, the number of mitochondria (n=179, *r*=0.56, *p*<10^−15^; Pearson correlation) and the total amount of mitochondria (n=179, *r*=0.68, *p*≈0; Pearson correlation) are also correlated with in-degree, the number of incoming synapses. (Figures 2B and S2B). These results suggest that neurons with more mitochondria tend to form more synaptic connections.

**Figure 2.**
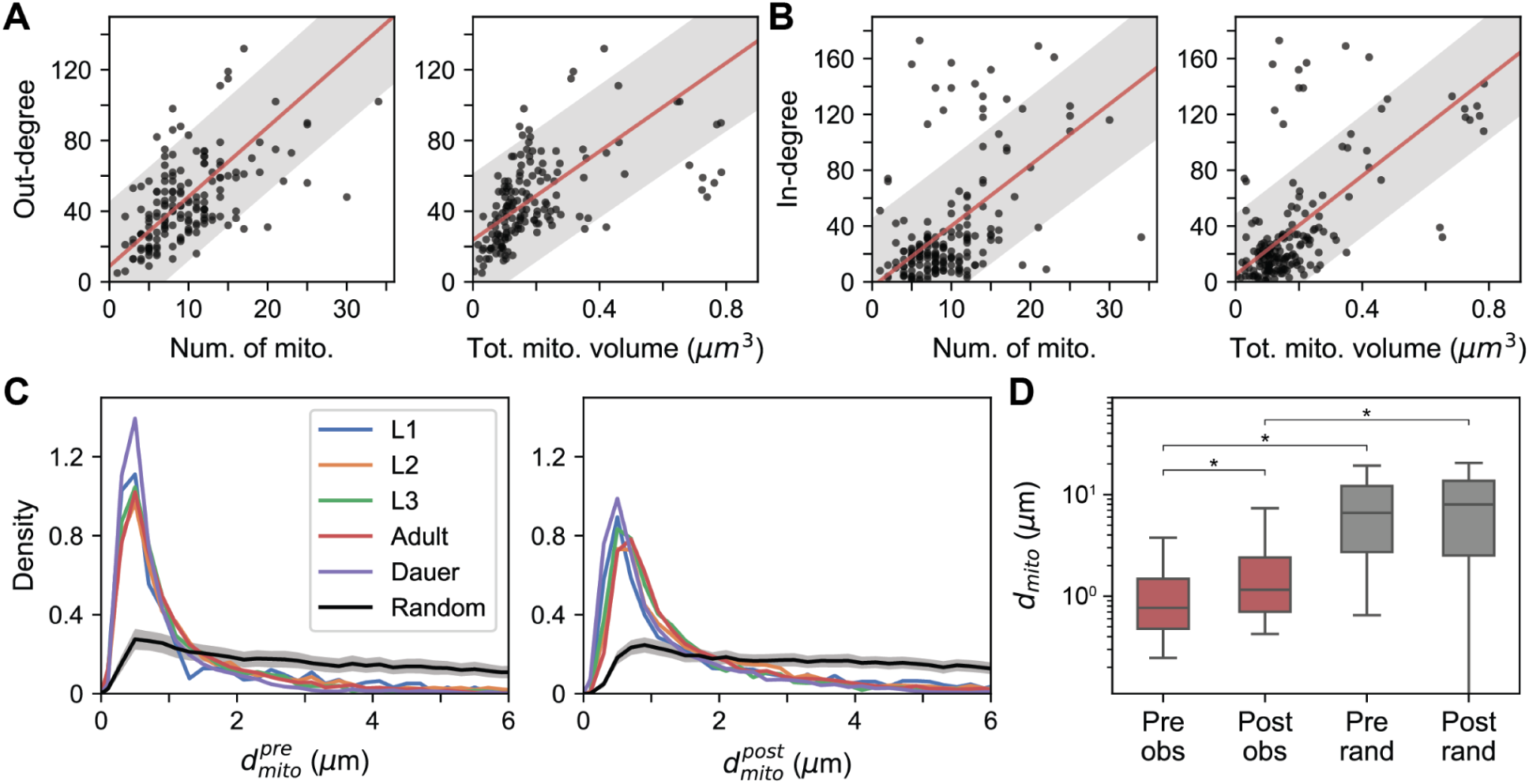
Mitochondria location is related to synaptic connections. (A) Out-degree is positively correlated with number of mitochondria in neuron (left: n=161, *r*=0.65, *p*=1.70×10^−20^; Pearson correlation) and sum of the mitochondria volume in neuron (right: n=161, *r*=0.62, *p*=2.46×10^−18^; Pearson correlation). RIAs are not shown for visualization purposes as they have outlying out-degree (see Figure S2). (B) Same with (A) for in-degree (left: n=179, *r*=0.56, *p*=1.89×10^−16^; right: n=179, *r*=0.68, *p*≈0; Pearson correlation). (C) Distance to nearest mitochondria from presynaptic (left) and postsynaptic (right) sites in different developmental stages (color lines) and distance to randomly assigned mitochondria (black line). Area under the curve is normalized to be equal to 1. (D) Distance to postsynaptic sites is greater than distance to presynaptic sites (n_pre_=3677, n_post_=7099, *p*≈0) and distance to randomly assigned mitochondria is significantly larger than distance to nearest mitochondria (*p*≈0). (A and B) Line: linear fit, shade: 80% prediction interval. (C) Black line: mean, black shade: 95% confidence interval. (D) Center line: median, box: interquartile range, whiskers: 5th and 95th percentile. \**p*≈0; Wilcoxon rank-sum test. (A-D) Adult data are shown as representative. See also Figure S2.

Nevertheless, the correlation between mitochondria morphology and out-degree does not explain how mitochondria and synapses are spatially correlated, as mitochondria might be concentrated at certain locations in neurites regardless of synapse locations. Therefore, we measured the distance from each active zone 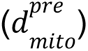 and each postsynaptic site 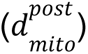 to the nearest mitochondria. Looking at the distribution of these distances, we were able to observe a peak within 1 µm in every developmental stage (Figure 2C). Considering the diameter of a bouton in the neurite is approximately in the range of 1 to 2 µm, this means there are mitochondria located at the same bouton where there are outgoing or incoming synapses. Consistent with previous results in *Drosophila*, we found that distances to postsynaptic sites are significantly larger than distances to active zones (Figures 2D and S2C; Rivlin et al. 2024; Riboul et al. 2023). To test this result did not occur by chance, we have assigned a random mitochondrion within a neuron for every active zone (Methods) and measured the distance to the nearest mitochondria. We were able to see that the distribution of distances in the randomized configurations were relatively more uniform and the distances in our data were significantly smaller in the observed configuration for both pre– and postsynaptic sites (Figures 2C and 2D; n_pre_=3677, n_post_=7099, *p*≈0; Wilcoxon rank-sum test).

Mitochondria support synaptic transmission by regulating neurotransmitter release by several mechanisms like calcium buffering. Therefore, mitochondria proximity to synapses could be important and we hypothesized mitochondria would be closely located to support larger synapses (i.e. larger active zones). Indeed, we found that large synapses tend to have the nearest mitochondria closer than smaller synapses (Figures 3A and S3A). Most of the synapses, over 95%, had similar synapse size, less than 0.001 µm^3^ but there were large synapses which have a diverse range of sizes (Figure 3A). Synapses that have mitochondria nearby (Methods) were significantly larger than synapses with mitochondria farther away (Figure 3A; n_near_=2408, n_far_=1269, *p*<10^−11^; Wilcoxon rank-sum test). This result is consistent with study in mammalian neurons (Smith et al. 2016), providing additional evidence for the relationship between mitochondria proximity and synapse efficacy. Vice versa, we measured the distance to pre-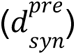 and postsynaptic 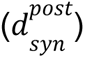 from the nearest mitochondria and observed mitochondria located closer to synapses were larger, indicating larger mitochondria are needed to operate larger synapses (Figures 3B, S3B, and S3C; pre: n_near_=1274, n_far_=219, *p*<10^−4^, post: n_near_=1249, n_far_=465, *p*<10^−5^; Wilcoxon rank-sum test).

**Figure 3.**
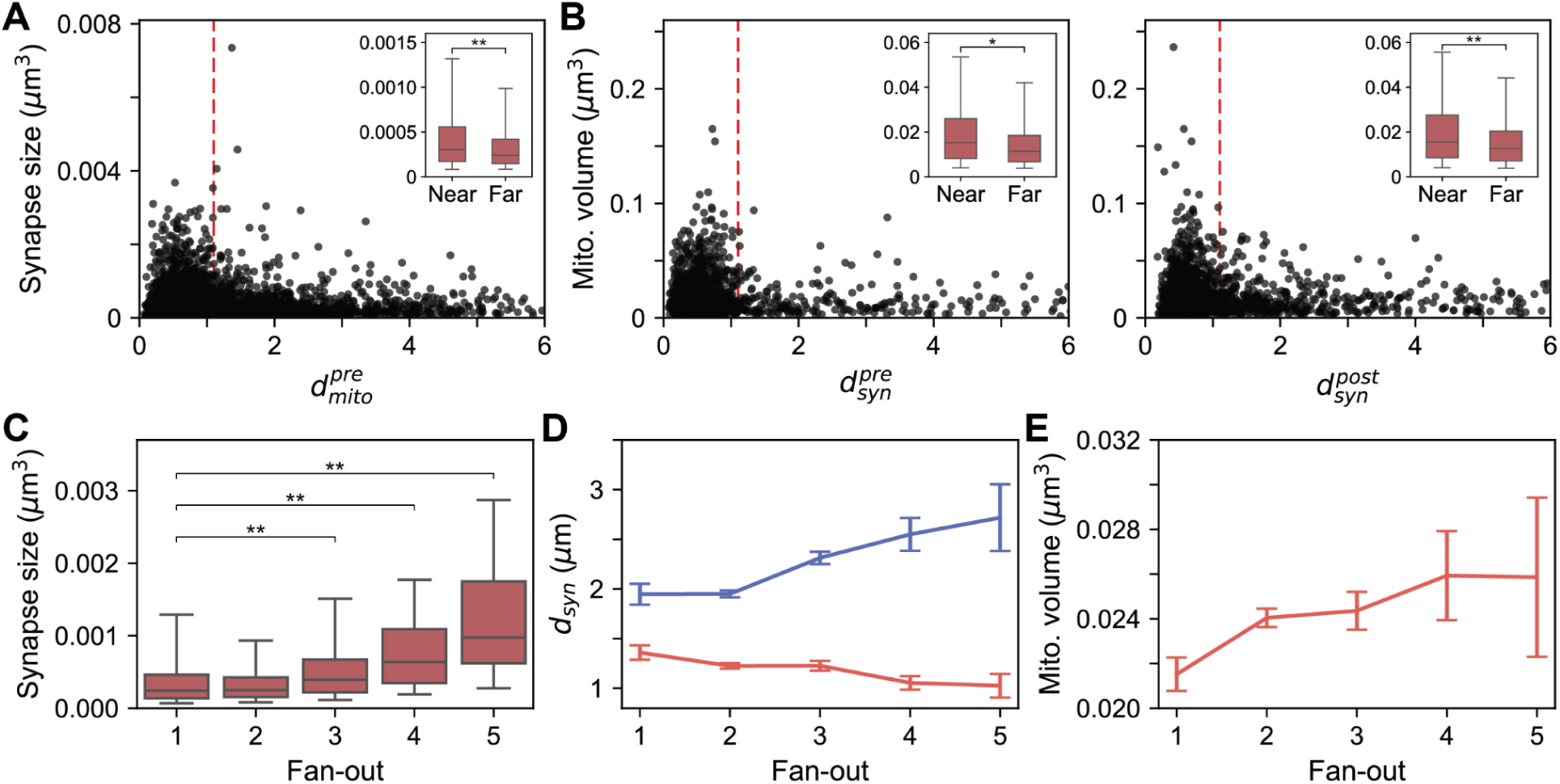
Mitochondria size is related to synaptic connections. (A) Active zones near (below red dashed) mitochondria are larger (inset) than those farther away (n_near_=2408, n_far_=1269, *p*=3.96×10^−12^). (B) Mitochondria near (below red dashed) synapses are larger (inset) than those farther away for both pre-(left; n_near_=1274, n_far_=219, *p*=6.16×10^−5^) and postsynaptic sites (right; n_near_=1249, n_far_=465, *p*=1.19×10^−6^). (C) Active zone sizes tend to be larger for synapses with larger fan-out (from left; *p*=2.83×10^−16^, *p*=2.48×10^−20^, *p*=2.92×10^−13^). (D) Mitochondria are located closer to presynaptic sites (red) and farther away from postsynaptic sites (blue) for synapses with larger fan-out. (E) Mitochondria near presynaptic sites are larger for synapses with larger fan-out. (A-C) Center line: median, box: interquartile range, whiskers: 5th and 95th percentile. \**p*<10^−4^, \*\**p*<10^−5^; Wilcoxon rank-sum test. (C-E) Pre: n_1_=514, n_2_=2303, n_3_=698, n_4_=125, n_5_=35; Post: n_1_=500, n_2_=4383, n_3_=1785, n_4_=361, n_5_=66. (A-E) Adult data are shown as representative. See also Figures S3 and S4.

*C*. *elegans* is a polyadic animal, meaning a single active zone can send neurotransmitters to multiple partners. From above results, we hypothesized that active zones with higher fan-out, number of postsynaptic partners per active zone, would have larger mitochondrion nearby as active zones with larger fanouts have larger synapses (Figures 3C and S4A). As expected, the nearest mitochondria were larger and closer to the synapses as fan-out increased (Figures 3D, 3E, S4B, and S4C). On the contrary, nearest mitochondria were farther away at the postsynaptic sites when they shared input with more cells (Figures 3D and S4B). All the relations were consistent among different developmental stages (Figures S2-S4).

### Axonal mitochondria are shorter than dendritic mitochondria

Mitochondria morphologies differ in different compartments (e.g. axon, dendrite, and soma) in mammalian neurons (D. T. W. Chang, Honick, and Reynolds 2006; Lewis et al. 2018b; Turner et al. 2022) as each compartment serves a different function. In *C*. *elegans* neurons, roles of different compartments are merged as the same neurite can both send and receive signals. However, there are exceptions in symmetric motor neurons. For example, it is known that SMDD and SMDV have outgoing synapses only at boutons in dorsal and ventral regions of arbor (Figures 4A-4C and S5A-S5D) respectively, which leads to compartmentalized activity in RIA (Hendricks et al. 2012). Unlike outgoing synapses, the incoming synapses exist at both boutons (Figures 4A-4C). Other classes of motor neurons such as RMD and SMBD neurons also show compartmentalized synapse distribution (Figures 4E, 4F, and S6A-S6D). A similar pattern is observed in RIA neurons as well (Figures S6E and S6F).

**Figure 4.**
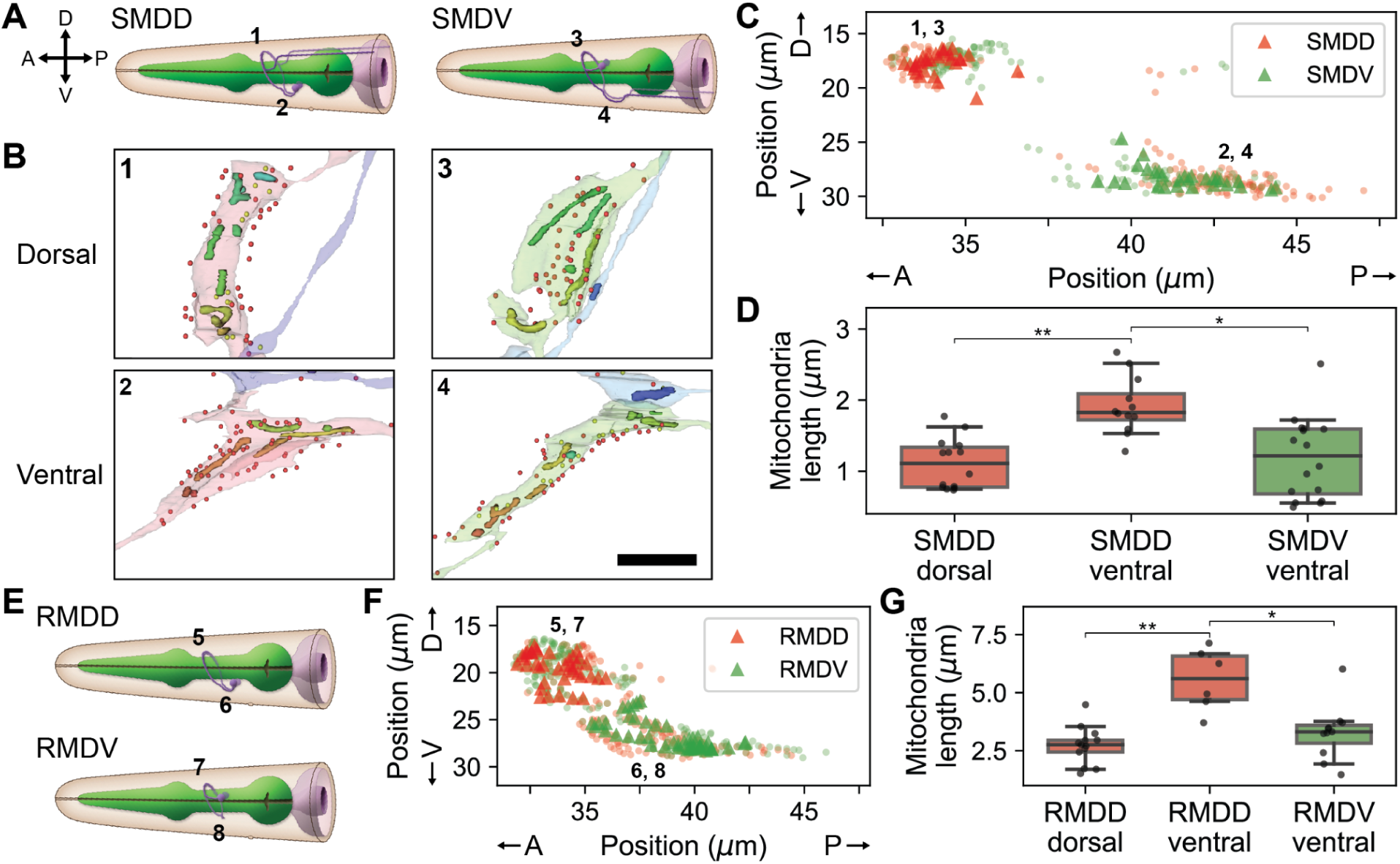
Axonal mitochondria are shorter. (A) Diagram of SMDD (left) and SMDV (right) neurons. (B) Reconstructed mitochondria in adult SMDD (left) and SMDV dorsal (top) and ventral (bottom) boutons. Number indicates locations marked in (A). Presynaptic (yellow) and postsynaptic (red) sites are marked with dots. (C) Distribution of outgoing (triangle) and incoming (circle) synapses of SMDD (red) and SMDV (green). Numbers are locations of numbered boxes in (B). (D) Mitochondria are longer in the ventral bouton of SMDD, where it only receives inputs (left, n_d_=14, n_v_=12, *p*=0.00011). Within the ventral boutons, mitochondria are longer in SMDD (right, n_SMDD_=12, n_SMDV_=16, *p*=0.00116). (E) Diagram of RMDD (top) and RMDV (bottom) neurons. (F) Same with (C) for RMDD (red) and RMDV (green). (G) Mitochondria are longer in the ventral bouton of RMDD, where it only receives inputs (left, *p*=0.00086). Within the ventral boutons, mitochondria are longer in RMDD (right, *p*=0.00356). (A and E) Diagrams are adopted from WormAtlas (Altun, Z.F. and Hall, D.H. 2024). A: anterior, P: posterior, D: dorsal, V: ventral. (B) Scale bar: 2 µm. (D and G) Center line: median, box: interquartile range, whiskers: 5th and 95th percentile. \**p*<0.01, \*\**p*<0.001; Wilcoxon rank-sum test. See also Figures S5 and S6.

Due to compartmentalized synapse distribution, it is feasible to isolate the neurite region with only incoming synapses, like dendrites, and study whether there is any difference in mitochondria structures. Here, we will define mitochondria in this region with only incoming synapses as “dendritic” and the ones in the other regions as “axonal” for convenience even though the boutons have mixed pre– and postsynaptic sites. We discovered that dendritic mitochondria are longer compared to axonal mitochondria in SMD neurons (Figures 4B and S5). We have verified this by comparing mitochondria in SMDD dorsal (axonal) and SMDD ventral (dendritic) boutons (Figure 4D; n_d_=14, n_v_=12, *p*=0.00011; Wilcoxon rank-sum test). As this result could be due to the difference in the bouton size, when we compared the ventral boutons of SMDD and SMDV, the mitochondria in SMDV ventral (axonal) boutons were smaller than mitochondria in SMDD ventral (dendritic) boutons (Figure 4D; n_SMDD_=12, n_SMDV_=16, *p*=0.00116; Wilcoxon rank-sum test). Similar results have been tested with RMD neurons and found mitochondria in RMDD ventral boutons are longer than those in RMDD dorsal (n_d_=13, n_v_=6, *p*=0.00086; Wilcoxon rank-sum test) and RMDV ventral boutons (Figure 4G; n_SMDD_=6, n_SMDV_=11, *p*=0.00356; Wilcoxon rank-sum test). This result is consistent with previous findings that dendritic mitochondria tend to be larger and longer (D. T. W. Chang, Honick, and Reynolds 2006; Lewis et al. 2018b; Turner et al. 2022). This structural property is a fundamental principle that can be found in all developmental stages (Figure S5).

### Mitochondria morphology is adapted to accommodate neural circuit function

Above, we have seen that axonal mitochondria have shorter morphology and mitochondria are spatially related to synapses. Therefore, we questioned whether the particular mitochondria morphology and localization is necessary for proper synaptic transmission, thereby important for intended behavioral output. SMD neurons have been known to be involved in omega turns or sharp directional changes as they activate head and neck muscles (White et al. 1986; Gray, Hill, and Bargmann 2005). Assuming mitochondria morphology is adapted to support synaptic transmission, we hypothesized disruption of mitochondria structures in SMD neurons would interrupt innervation of body wall muscles, inhibiting the turning behavior.

In this study, we studied a kind of exploratory behavior, local search, to investigate how mitochondria structure affects the behavior (Figure 5A; Gray, Hill, and Bargmann 2005). When the animals are removed from food, they make sharp turns to search for food, mediated by SMD neurons, similar to biased random walk (Pierce-Shimomura, Morse, and Lockery 1999; Gray, Hill, and Bargmann 2005). As it has been previously reported, animals showed high turning rate due to local search behavior immediately after they have been transferred to an environment without food (Figure 5B). Then the turning rate relaxed to the base rate where they stopped showing local search behavior (Figure 5B). According to our hypothesis, we expected this local search behavior would be impaired when the mitochondria structure is disrupted.

**Figure 5.**
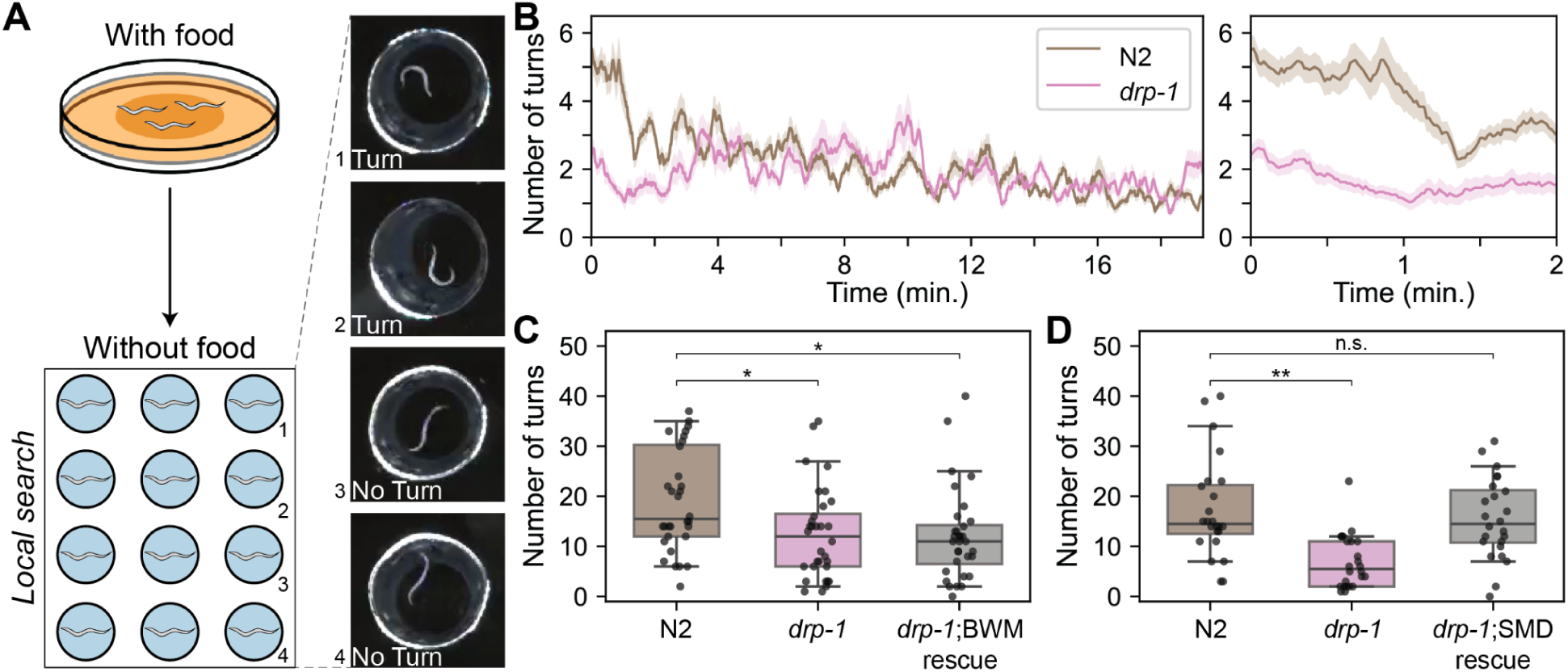
Mitochondria morphology is adapted to accommodate neural circuit function. (A) Behavior experiment setup. Worms were transferred from the environment with food to the environment without food, then the number of turns were counted. (B) Number of turns over time for 20-minute recording (left) and for first 2-minute recording (right). (C) Total number of turns in the first 2 minutes for N2 wild-type, *drp-1* mutants, and BWM-specific rescue model. *drp-1* mutants exhibit significantly lower number of turns compared to the wild-type and impaired behavior in *drp-1* mutants are not recovered in the BWM-specific rescue model (n=32, *p*_drp1_=0.009, *p*_rescue_=0.003). (D) Same with (C) for SMD neurons and the behavior is recovered in the SMD-specific rescue model (n=24, *p*_drp1_=1.64×10^−5^, *p*_rescue_=0.66). (C and D) Center line: median, box: interquartile range, whiskers: 5th and 95th percentile. \**p*<0.01, \*\**p*<0.0001; Wilcoxon rank-sum test.

To test this idea, we checked the local search behavior in the mutants important for the mitochondria morphology *drp-1*, a mitochondria fission factor. In fact, *drp-1* mutants, where mitochondria fission is interrupted, showed a defect in increased turning rate during local search behavior (Figure 5B). This result suggests that mitochondria morphology and localization regulated by the mitochondria fission factor is important for proper synaptic function in SMD neurons.

To quantitatively compare the local search behavior, we counted the total number of turns within the first 2 minutes after the worms were transferred, where an increased turning rate was observed in wild-type animals (Figure 5B). The total number of turns were significantly lower in *drp-1* mutants compared to the wild-type as expected (Figures 5C and 5D; n_N2_=32, 24, n_drp1_=32, 24, *p*=0.009, *p*=1.64×10^−5^; Wilcoxon rank-sum test). However, *drp-1* mutants have broken mitochondria structure in various cells. We first tested whether this deficit in behavior is caused by the disruption of mitochondria morphology in muscle cells as they are the cells that execute the behavior. However, the total number of turns in BWM-specific rescue of *drp-1* mutants were not significantly different from that of *drp-1* mutants (Figure 5C; n_N2_=32, n_rescue_=32, *p*=0.003; Wilcoxon rank-sum test).

With BWMs excluded from the candidates, we then moved onto our original hypothesis, that this behavior deficit is due to the dysfunction in SMD neurons, which fails to innervate BWMs. We created SMD-specific rescue of *drp-1* mutants and were able to see the local search behavior recovered to the normal level (Figure 5D; n_N2_=24, n_rescue_=24, *p*=0.66). These results imply mitochondria structure is adapted to accommodate proper synaptic transmission in neurons to maintain effective functioning of the neural circuit.

### Dauer inter– and motor neurons show distinctive mitochondrial features

Beyond the fundamental structural properties of mitochondria conserved across development, are there any differences? Since we have densely reconstructed mitochondria in diverse developmental stages, it enables us to conduct stage-wise quantitative comparison of mitochondria structure. As mitochondria structure is related to synaptic connections (Figures 2 and 3), investigating mitochondria morphologies of neurons in different stages could provide insights in understanding the differences in neural circuits.

We have computed various mitochondrial features including total mitochondria volume, surface area, volume fraction, length, and mitochondrial complexity index (MCI; Vincent et al. 2019), Methods). We initially explored an overview of mitochondrial features by projecting all mitochondria features onto principal components. As a result, we observed that several dauer neurons are separately clustered from the majority, indicating that neuronal mitochondria in dauer are distinctive (Figures 6A and S7A).

**Figure 6.**
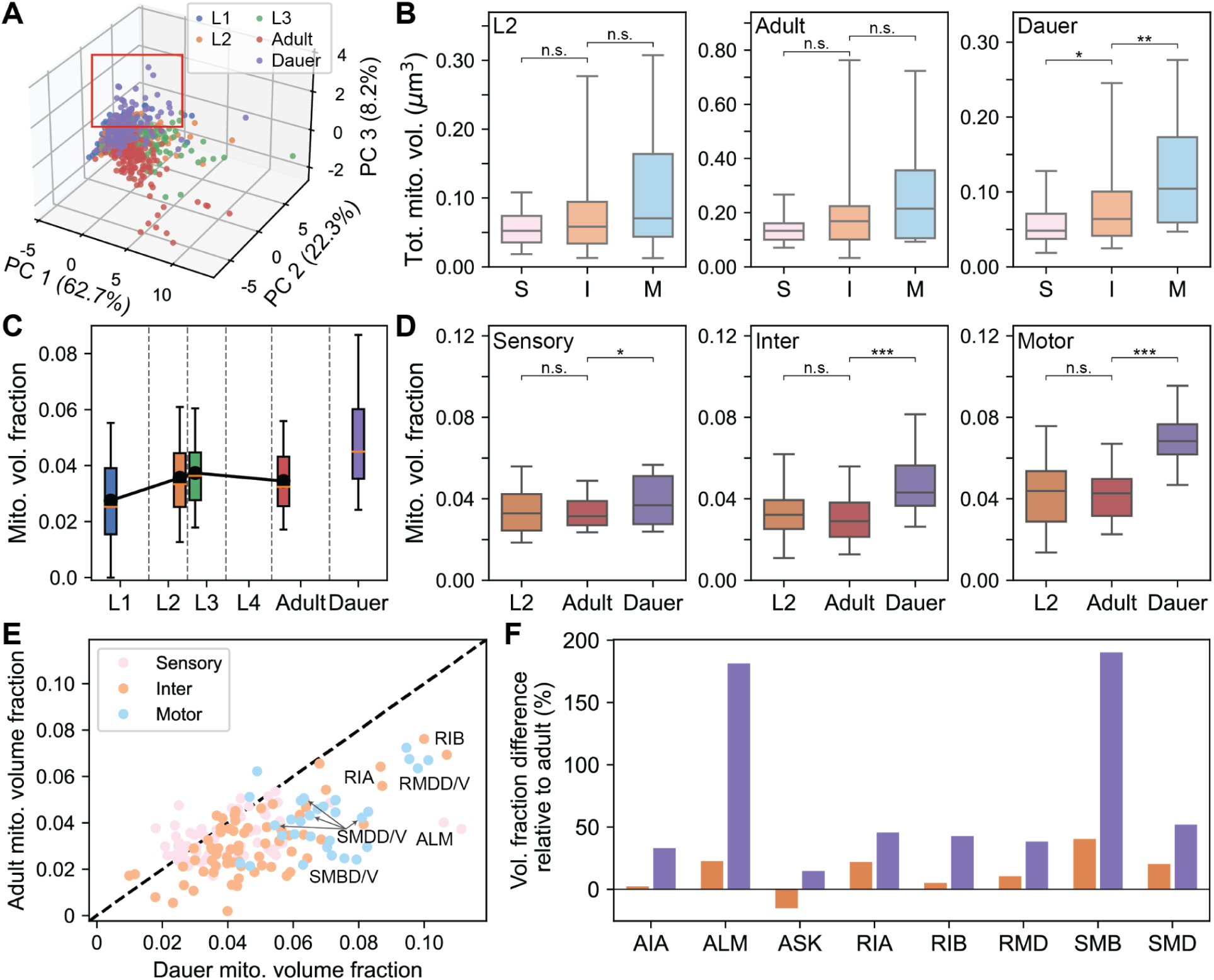
Dauer inter– and motor neurons show distinctive mitochondrial features. (A) Principal component embeddings of neuronal mitochondrial features show group of dauer neurons clustered outside (red box). (B) Distribution of total mitochondria volume in neurons per type for L2 (left; *p*=0.35, *p*=0.15), adult (middle; *p*=0.12, *p*=0.09), and dauer (right; *p*=0.014, *p*=0.004). (C) Distribution of mitochondria volume fraction across development. (D) Distribution of mitochondria volume fraction in L2, adult, and dauer for sensory (left; *p*=0.772, *p*=0.045), inter-(middle; *p*=0.333, *p*=2.99×10^−8^), and motor (right; *p*=0.778, *p*=3.69×10^−8^) neurons. (E) Comparison between mitochondria volume fraction of dauer and adult. Dauer shows higher mitochondria volume fraction relative to other developmental stages. Dashed line indicates *y* = *x* line. (F) Difference in mitochondria volume fraction compared to selected neurons in adult for L2 (orange) and dauer (purple). Volume fraction difference is larger in the dauer stage. (C) Black: mean. (B-D) Center line: median, box: interquartile range, whiskers: 5th and 95th percentile. (B and D) n_sen_=61, n_int_=69, n_mot_=32, \**p*<0.05, \*\**p*<0.01, \*\*\**p*<10^−6^, \*\*\*\**p*<10^−7^; Wilcoxon rank-sum test. See also Figure S7.

Neurons in *C*. *elegans* are classified into three types – sensory, inter-, and motor neurons – depending on their functions. Variability in function can be encoded in their arbors as the cell shapes of different types differ and mitochondria structure can also differ accordingly. In most of the stages, no significant difference in the distribution of total mitochondria volume was observed between different cell types, even though there were some interneurons and motor neurons that possess large amounts of mitochondria (Figures 6B and S7B). On the other hand, motor neurons in dauer had greater total mitochondria volume on average compared to interneurons and sensory neurons (Figure 6B; n_sen_=61, n_int_=69, n_mot_=32, *p*=0.004; Wilcoxon rank-sum test). Moreover, interneurons in dauer had significantly more mitochondria than sensory neurons (Figure 6B; *p*=0.014; Wilcoxon rank-sum test).

Still, we lack evidence to draw conclusions regarding stage-wise comparisons because size of neurons differ between stages and it is obvious that larger cells are likely to have more mitochondria (Figures S1E-S1G). In order to compensate for this caveat, we introduced a measure, mitochondria volume fraction, where we divide the total mitochondria volume in a neuron by the neuron arbor volume. Unlike the mitochondria volume, mitochondria volume fraction was constant in normal reproductive stages, from L1 to adult stages except for the dauer stage, where the mitochondria volume fraction was exceptionally high (Figure 6C). When we analyze per cell types, the mitochondria volume fraction of neurons in dauer were significantly greater in interneurons (n_int_=69, *p*=0.333, *p*=2.99×10^−8^; Wilcoxon rank-sum test) and motor neurons (n_mot_=32, *p*=0.778, *p*=3.69×10^−8^) while the difference was negligible in sensory neurons (Figure 6D; n_sen_=61, *p*=0.772, *p*=0.045; Wilcoxon rank-sum test). As the amount of mitochondria is correlated with number of connections, this result could imply increased role of motor neurons in neural circuits of dauer compared to normal reproductive stages, which is consistent with recent findings in *C*. *elegans* dauer connectome (Yim et al. 2024).

To verify that this difference of mitochondria volume fraction is true for the same neuron in different stages, we looked at the mitochondria volume fraction of individual neurons (Figures 6E). Mitochondria volume fraction of neurons in dauer were substantially larger for many neurons (Figures 6E and 6F) while volume fraction of neurons in other stages were similar (Figures S7D and S7E). As we saw from the distributions, many interneurons and motor neurons tend to have higher mitochondria volume fraction, especially neurons involved in head and neck movement (Gray, Hill, and Bargmann 2005)Figures 6E and 6F; Gray, Hill, and Bargmann 2005).

### Dauer cholinergic and glutamatergic neurons show distinctive mitochondrial features

Given the critical role of mitochondrial function in neurotransmitter release, it is reasonable to assume that mitochondria structure may vary depending on neurotransmitter identities. Besides neuronal types, neurons can be classified based on the neurotransmitters they release, with possibility of diverse neurotransmitters within the same type (Hobert, Glenwinkel, and White 2016). In *C*. *elegans*, most neurons release acetylcholine (ACh) and glutamate (Glu), with sensory and interneurons primarily consisting of cholinergic and glutamatergic types (Figure 7A). Additionally, a few sensory neurons release dopamine, while several interneurons release GABA (Figure 7A). For this study, we restricted our analysis on the three predominant neurotransmitters – ACh, Glu, and GABA – since neurons releasing other neurotransmitters are too few to draw reliable conclusions.

**Figure 7.**
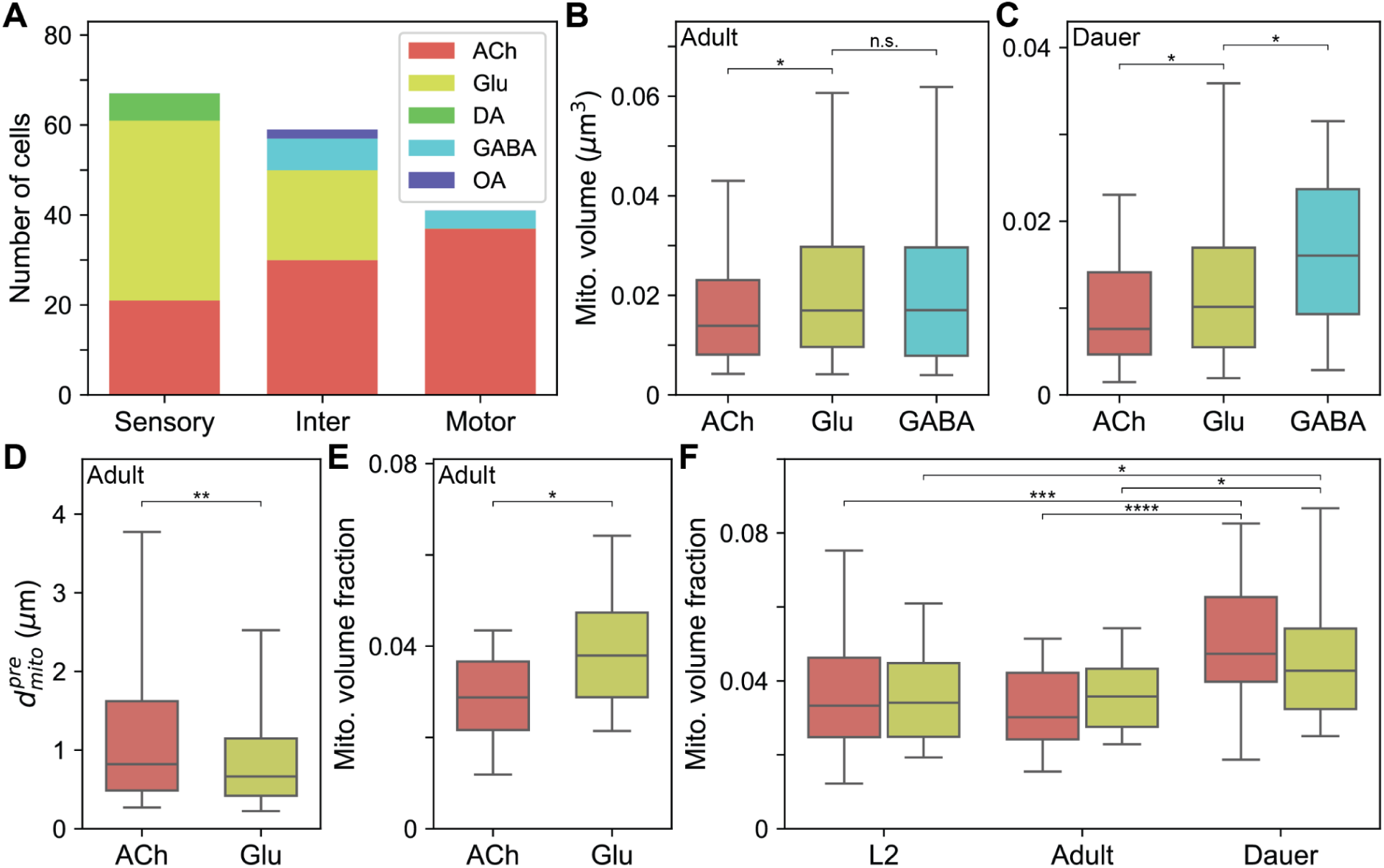
Dauer cholinergic and glutamatergic neurons show distinctive mitochondrial features. (A) Proportion of neuron types by neurotransmitter classification for each cell type. (B) Glutamatergic interneurons have larger mitochondria than cholinergic interneurons (n_ACh_=267, n_Glu_=308, n_GABA_=108, *p*=0.001, *p*=0.797). (C) Same with (B) but GABAergic interneurons have larger mitochondria than glutamatergic interneurons in dauer (n_ACh_=186, n_Glu_=199, n_GABA_=48, *p*=0.002, *p*=0.007). (D) Glutamatergic interneurons have mitochondria closer to active zones than cholinergic interneurons (n_ACh_=428, n_Glu_=845, *p*=4.45×10^−6^). (E) Glutamatergic interneurons have higher mitochondria volume fraction than cholinergic interneurons (n_ACh_=30, n_Glu_=20, *p*=0.007). (F) Dauer cholinergic (n=78, 85, 85, *p*=7.95×10^−7^, *p*=3.48×10^−9^) and glutamatergic neurons (n=58, *p*=0.003, *p*=0.003) have higher mitochondria volume fraction. (A-E) Ach: Acetylcholine, Glu: Glutamate, DA: Dopamine, OA: Octopamine. (B-F) Center line: median, box: interquartile range, whiskers: 5th and 95th percentile. \**p*<0.01, \*\**p*<10^−5^, \*\*\**p*<10^−6^, \*\*\*\**p*<10^−8^; Wilcoxon rank-sum test. See also Figure S8.

We focused our analysis on interneurons because these neurons primarily connect with other neurons rather than non-neuronal cells. Among interneurons, we found that mitochondria in glutamatergic and GABAergic neurons have larger mitochondria than those in cholinergic neurons in normal reproductive stages (Figures 7B and S8A; n_ACh_=267, n_Glu_=308, n_GABA_=108, *p*=0.001, *p*=0.797; Wilcoxon rank-sum test). Alternatively, GABAergic interneurons possess even larger mitochondria than glutamatergic interneurons in the dauer stage (Figure 7C; n_ACh_=186, n_Glu_=199, n_GABA_=48, *p*=0.002, *p*=0.007; Wilcoxon rank-sum test).

Since glutamatergic neurons have larger mitochondria on average, we hypothesized that mitochondria in these neurons might be located closer to synapses. We found that glutamatergic interneurons have the nearest mitochondria more closely located to the active zones (Figure 7D; n_ACh_=428, n_Glu_=845, *p*=4.45×10^−6^; Wilcoxon rank-sum test) and also have higher mitochondria volume fraction (Figure 7E; n_ACh_=30, n_Glu_=20, *p*=0.007; Wilcoxon rank-sum test) compared to cholinergic interneurons. These findings were also observed in other developmental stages (Figures S8B and S8C) and align with previous reports in *Drosophila* whole brain, suggesting this may be a shared characteristic among invertebrates (Rivlin et al. 2024).

From above results, it can be inferred that more mitochondria are needed closer to synapses in glutamatergic and GABAergic neurons (Figures 7B-7E). Considering GABA is inhibitory neurotransmitter and glutamate can act as inhibitory neurotransmitter in some neurons like AIB and RIM, this result could be indicating inhibitory interneurons require more energy per cell compared to other excitatory neurons (Kann 2016; Rivlin et al. 2024).

Although structural properties of mitochondria are generally consistent across development among neurons releasing the same neurotransmitter, variations were also noted between different stages. We noticed that both cholinergic (n_L2_=78, n_adult_=85, n_dauer_=85, *p*=7.95×10^−7^, *p*=3.48×10^−9^; Wilcoxon rank-sum test) and glutamatergic (n=58, *p*=0.003, *p*=0.003; Wilcoxon rank-sum test) neurons in dauer, regardless of the cell types, show higher mitochondria volume fraction than normal reproductive stages (Figure 7F). Acetylcholine and glutamate are known to play crucial roles during the dauer stage such as modulating behavioral responses specific to the dauer stage. The mitochondria structure could have been adapted to accommodate the stage-specific needs in cholinergic and glutamatergic neurons.

### Dauer body wall muscle mitochondria exhibit distinctive structure

We have not only reconstructed neuronal mitochondria but also reconstructed those in body wall muscles (BWMs), allowing us to make structural comparisons of BWM mitochondria across development. The BWMs included in the data are head and neck muscles as our study has been centered around the vicinity of the nerve ring.

Interestingly, the morphologies of mitochondria in head and neck muscles undergo substantial changes throughout the development (Figure 8A). In L1, muscle cells contain a solitary strand of mitochondria, and as the worm matures, it branches into additional mitochondria strands (Figure 8A). Once the worm becomes an adult, the mitochondria form a reticulum-like structure with many strands intermingled (Figure 8A). Similar developmental changes have been observed using fluorescence imaging in BWMs (Schultz et al. 2017; Han et al. 2012). In dauer, the BWMs exhibit a distinctive reticulum-like structure, which distinguishes from the adult stage, where the strands are slimmer and adjacent portions of the muscle belly contain greater empty spaces, occupied by lipid droplets (Figures 8A and S9A; Popham and Webster 1979). To confirm that our findings are not limited to specific animals, we acquired fluorescence images of mitochondria in head and neck muscles. We have verified that our observation is valid in different animals, further highlighting the distinctive reticulum-like structure in the dauer stage (Figure 8A).

**Figure 8.**
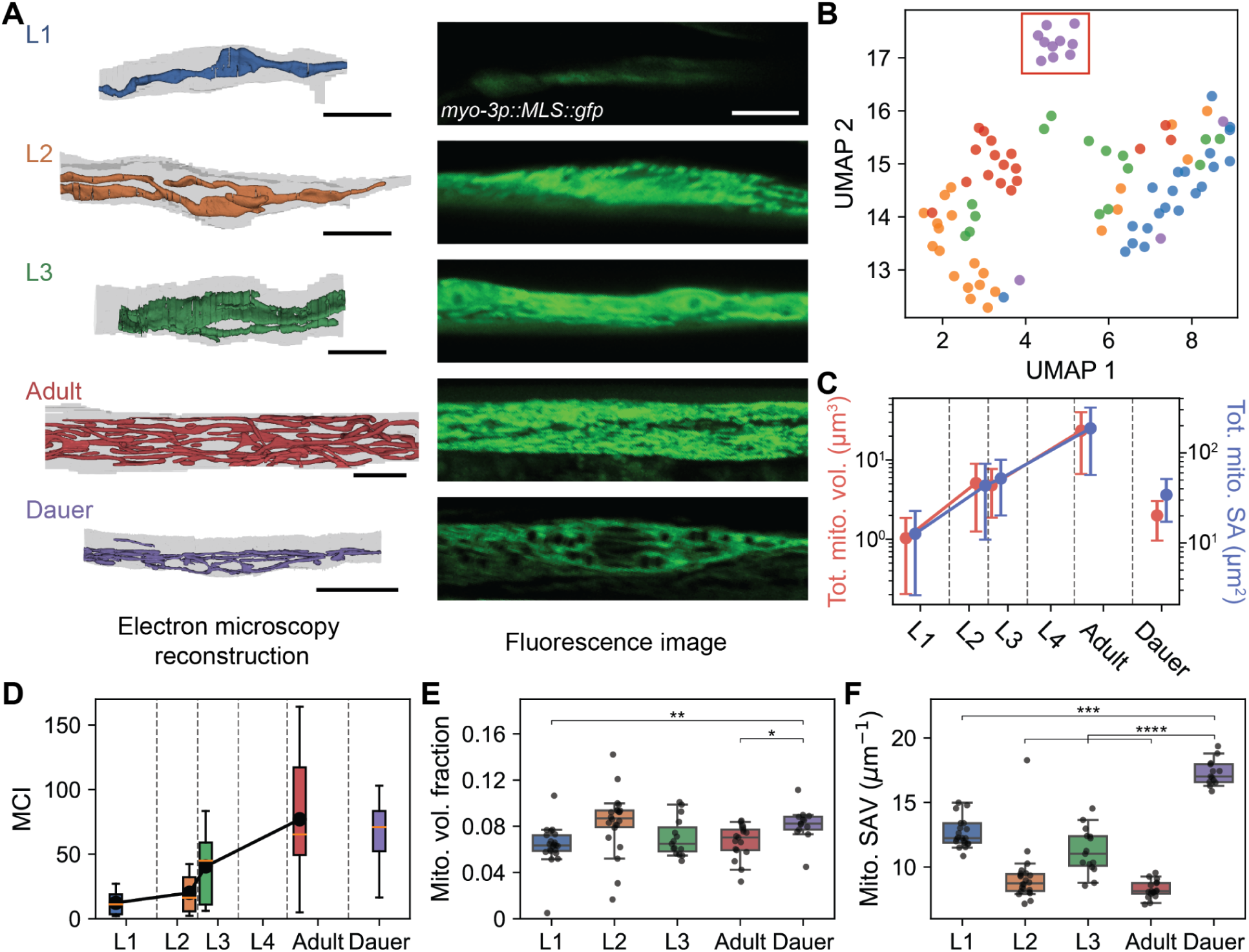
Dauer body wall muscle mitochondria exhibit distinctive structure. (A) Reconstructed mitochondria in example body wall muscles (BWMs) in different developmental stages (left) and corresponding fluorescences images of mitochondria in BWM (right). (B) UMAP embeddings of BWM mitochondria features show group of dauer cells clustered (red box). (C) Total mitochondria volume (red) and total mitochondria surface area (blue) across development (mean±std). (D) Mitochondria complexity index (MCI) across development. (E) Mitochondria volume fraction across development (*p*_L1_=4.22×10^−4^, *p*_L2_=0.34, *p*_L3_=0.05, *p*_adult_=0.003). (F) Mitochondria surface area per volume (SAV) across development (*p*_L1_=1.21×10^−5^, *p*_L2_=6.59×10^−6^, *p*_L3_=7.08×10^−6^, *p*_adult_=3.75×10^−6^). (A and E) Scale bars: 5 µm (black), 5 µm (white). (B-F) n_L1_=21, n_L2_=22, n_L3_=15, n_adult_=17, n_dauer_=13. (D-F) Center line: median, box: interquartile range, whiskers: 5th and 95th percentile. (E and F) \**p*<0.01, \*\**p*<0.001, \*\*\**p*<10^−4^, \*\*\*\**p*<10^−5^; Wilcoxon rank-sum test. See also Figure S9.

Since we have high-resolution reconstructions, we can also quantify the BWM mitochondria morphologies and quantitatively confirm the above results from qualitative investigation. When analyzing various mitochondrial features using UMAP embeddings, we found dauer muscle cells form a separate cluster, confirming that mitochondria in the dauer stage display distinctive characteristics (Figure 8B, Methods; McInnes, Healy, and Melville 2018). Upon closer examination, we noted an exponential increase in mitochondria volume and surface area from L1 stage to adulthood in BWMs (Figure 8C). Dauer BWMs had less total mitochondria volume compared to L2 and L3 likely due to enlarged sarcomeres and space occupied by lipid droplets, although mitochondria surface area remained similar (Figures 8C and S9A; Popham and Webster 1979; Wolkow and Hall 2016). The MCI also increased from L1 to adulthood, reflecting more complex, network-like mitochondria structure in adults (Figures 8A and 8D). Notably, dauer BWM mitochondria exhibited a high MCI due to its reticulum-like morphology (Figure 8D).

Given the reduced muscle belly in dauer BWMs, we hypothesized that the proportion of area occupied by mitochondria might be larger. However, the mitochondria volume fraction was not significantly different from other stages, with the exception of L1 (Figure 8E). Instead, the surface area per volume of mitochondria in dauer BWMs was significantly higher than in other stages (Figures 8C and 8F; n_L1_=21, n_L2_=22, n_L3_=15, n_adult_=17, n_dauer_=13, *p*_L1_=1.21×10^−5^, *p*_L2_=6.59×10^−6^, *p*_L3_=7.08×10^−6^, *p*_adult_=3.75×10^−6^; Wilcoxon rank-sum test). This result reflects the qualitative observation that dauer BWM mitochondria are composed of networks of thin strands (Figure 8A).

Next, we checked whether this dauer-specific mitochondria structure is preserved even after the dauer exit. Captured with fluorescence imaging, BWM mitochondria in postdauer L4 or postdauer adult stages did not show distinctive reticulum-like structure as seen in dauer (Figure S9B). Mitochondria in postdauer L4 and postdauer adult did not have empty spaces in between the strands which were filled with lipid droplets like dauer (Figure S9A).

## Discussion

In this study, we present comprehensive dense reconstructions of mitochondria in normal reproductive stages and the dauer stage of *C*. *elegans* using 3D electron microscopy. Utilizing these reconstructed datasets, we conducted a comparative analysis of mitochondria structures throughout development, spanning from birth to adulthood, in both neurons and body wall muscles. Consequently, we were able to propose fundamental mitochondria structural principles shared across different stages as well as stage-specific mitochondrial features. EM offers a distinct advantage due to its superior resolution, enabling detailed quantitative morphological comparisons that are often not feasible with fluorescence imaging, especially in small animal models such as *C*. *elegans*. Furthermore, our investigation encompassed the examination of mitochondria across the nervous system from sensory neurons to muscles. This provides insights into how mitochondria structure might impact neural circuit function leading to behavioral output. Lastly, we observed unique mitochondria structures in the alternative dauer stage, which nematodes enter under adverse environmental conditions, suggesting potential adaptation of intracellular organelles.

### Fundamental structural principles are preserved across development

We observed many structural properties of mitochondria are preserved throughout the development, including during the alternative dauer stage. The relationship between the quantity and proximity of mitochondria to the synaptic connectivity demonstrates consistent rules across all stages (Figures 2, 3, S2-S4). Similarly, compartment-specific mitochondria morphology is conserved across developmental stages (Figures 4, S5, and S6). In addition, mitochondria exhibit comparable characteristics in each cell type during normal reproductive stages (Figures 6 and S7), even when cells are classified by the neurotransmitters they release (Figures 7 and S8). Cells and neural circuits have essential functions necessary for basic operational capabilities. Therefore, mitochondria need to follow established principles to support these essential functions.

These principles have been similarly identified in mammalian neurons. Previous studies have found the spatial distribution of synapses and mitochondria along the neurite covary, meaning synapses have mitochondria nearby potentially as an energy source and calcium buffer (Smith et al. 2016; Turner et al. 2022). Other studies have found that mitochondria in different compartments (i.e. axon, dendrites, soma) have different morphological characteristics. It has been discovered that dendritic mitochondria tend to be longer and larger than axonal mitochondria (Diane T. W. Chang, Honick, and Reynolds 2006; Lewis et al. 2018a; Turner et al. 2022). We now report similar findings in *C*. *elegans* indicating the same structural principle applies even in invertebrates, further supported by a recent comprehensive study of mitochondria in *Drosophila* (Rivlin et al. 2024). This provides a link between higher order animals and *C*. *elegans*, suggesting *C*. *elegans* as a great model organism to study how mitochondria structure affects synaptic function.

### Increased mitochondria in the dauer neurons for dauer-specific behavior

The notable increase in mitochondria volume fraction in dauer seems ironic since the worms in this stage display long periods of immotility, but with occasional bursts of fast actions (Cassada and Russell 1975; Gems et al. 1998). It is possible that dauer neurons contain more mitochondria relative to cell size as a preparatory mechanism when the occasion arises. The neurons that stand out with higher mitochondria volume fraction in dauer are interneurons (e.g., RIA, RIB) and motor neurons (e.g., RMD, SMD, SMB) that are known to be involved in head and neck neuromuscular system (Figures 6E and 6F). These cells are likely to be the drivers of forward and backward body movements. For instance, SMB neurons determine the amplitude of sinusoidal motion (Gray, Hill, and Bargmann 2005) and RMD neurons are known to control head-withdrawal reflex and spontaneous foraging behavior (Hart, Sims, and Kaplan 1995). A higher mitochondria density in these cells could enable neurons to rapidly activate synaptic connections, leading to quicker responses to the stimuli in the dauer stage.

Based on the previous observations, the dauer-specific behavior, nictation, is initiated from the head movement (Cassada and Russell 1975; H. Lee et al. 2011). The sensory neuron which regulates nictation, IL2, has been found (H. Lee et al. 2011), but the rest of the neurons in the neural circuit that generate nictation behavior are still unknown despite recent efforts in dauer connectome (Yim et al. 2024). Neurons like RIA, RIB, and SMD, which exhibit distinctive mitochondrial features, could be plausible candidates responsible for nictation (Figures 6E and 6F).

RMD, SMD, and IL2 neurons are all cholinergic, and given acetylcholine’s crucial role in the neuromuscular function, it is expected to play an important role during the dauer stage. Glutamatergic neurons, including RIB, are known to modulate behaviors in dauer (Zou et al. 2018). The finding that both cholinergic and glutamatergic neurons exhibit a high mitochondria volume fraction could be reflecting their important roles in the dauer stage (Figure 7F). Therefore, the neurons with higher mitochondria volume fraction in dauer could be candidates that regulate dauer-specific behavior, which can be tested with an optogenetics experiment (Yim et al. 2024).

Synaptic transmission not only influences animal behavior but also affects developmental decision making. Previous study reported that pheromone activates ASK and ADL neurons, which in turn stimulates AIA neurons via glutamatergic transmission to induce dauer entry (Chai et al. 2022). ASK neurons have high mitochondria volume fraction in dauer as other glutamatergic neurons (Figure 6F). Also, AIA neurons are interneurons that show high mitochondria volume fraction in dauer (Figure 6F), indicating the potential adaptation of mitochondria structure in the dauer stage.

### Mitochondria and lipid droplets in the dauer muscle cells

Previous studies using 2D cross-section analysis have shown that dauer muscle mitochondria are present in a compact conformation concentrated in the muscle belly (Popham and Webster 1979). It is presumed that this structural property is necessary to provide large amounts of energy on a short-term basis to support rapid response behavior (Cassada and Russell 1975). This study has not only confirmed that discovery but also extended it by uncovering the 3D structural morphology, including the sparse network-like structure with thin strands (Figure 8A).

Then the question naturally arises, what is present within the gaps amidst the strands? It has been reported that dauer muscle cells contain lipids unlike other developmental stages (Popham and Webster 1979). Indeed, the empty spaces between the mitochondria strands were occupied with lipid droplets forming intimate contacts with mitochondria (Figure S9A).

During the dauer stage, *C*. *elegans* transitions from glycolysis to using the glyoxylate cycle within mitochondria for energy production, enabling the conversion of stored fats into glucose (Burnell et al. 2005). The unique reticulum-like structure of BWM mitochondria observed in dauer (Figures 8A) may facilitate this metabolic shift. This could be a strategy adopted to increase the contact area between mitochondria and lipids to accommodate the efficient energy supply for the rapid and powerful response in dauer (Cassada and Russell 1975; O’Riordan and Burnell 1990). This result is consistent with previous findings in human skeletal muscles which showed the amount of lipid droplet and the contact area between lipids and mitochondria increased after intense exercise (Tarnopolsky et al. 2007).

### Generalizable mitochondria reconstruction pipeline

We developed an automated method for detecting mitochondria in EM, allowing us to acquire dense mitochondria reconstructions across various developmental stages. Initially, we trained our detection model with the dauer EM data and subsequently enhanced the model with additional ground truth from each dataset (Methods). Still, the initial model showed decent performance without fine-tuning, as mitochondria look similar in different EM datasets. Our final model, trained with all five datasets used in this study, demonstrates robust generalizability across new *C*. *elegans* EM datasets and potentially those from other animal models. Moreover, our model operates in 2D, making it accessible for researchers who work with single-section images to apply our method. This method could be extended to quantitatively analyze mitochondria morphology in disease models like Huntington’s disease or Alzheimer’s disease, where mitochondria dysfunction is prominent (Swerdlow 2011; Lopez Sanchez, van Wijngaarden, and Trounce 2019; Zhang et al. 2016; A. Lee et al. 2018; Liu et al. 2022; Neueder et al. 2024). All codes and models are publicly available at the GitHub repository (Methods).

## Acknowledgments

We thank Seok-Kyu Kwon and Yusuke Hirabayashi for valuable feedback and suggestions. We would like to thank Mei Zhen and her group for discussion and suggestions. Also, we thank the groups of Mei Zhen, Aravi Samuel, and Jeff Lichtman for generating and sharing non-dauer datasets that were used in this study. We are grateful to WormAtlas for providing invaluable resources and reference illustrations. The work was supported by the Samsung Science and Technology Foundation (SSTF-BA1501-52). J.A.B. acknowledges support by the National Research Foundation of Korea (NRF) grant (2019R1A6A1A10073437) funded by the Korean Ministry of Education. K.C.N. and D.H.H. were funded by NIH OD010943.

## Author Contributions

K.C.N and D.H.H. acquired dauer EM image stack. G.K., H.Y., and D.T.C. refined the cell segmentation of dauer dataset based on the initial segmentation generated by H.K. J.A.B. trained the mitochondria detection models and reconstructed mitochondria using the ground truth annotated by G.K. J.A.B. performed computational analysis. M.C. designed and implemented the microdroplet swimming assay and performed behavior experiments. S.A. acquired fluorescence images of body wall muscle mitochondria. S.A., M.C., D.T.C. acquired fluorescence images of neural mitochondria. J.A.B. wrote the paper with contributions from M.C., D.H.H., J.S.K., and J.L. J.A.B., J.S.K., D.H.H., and J.L. managed the multi-institution collaboration.

## Inclusion and Diversity

We support inclusive, diverse, and equitable conduct of research.

## Methods

### Electron microscopy images

For normal reproductive stages, we used electron microscopy (EM) images published by Witviet et al. (Witvliet et al. 2021) (L1: Dataset 2, L2: Dataset 5, L3: Dataset 6, Adult: Dataset 8). The raw images were acquired from Brain Observatory Storage Service and Database (BossDB) using *intern* library (Hider et al. 2022; Matelsky et al. 2021). For dauer stage, the same EM dataset from Yim et al. (Yim et al. 2024) was used. Please refer to Yim et al. (Yim et al. 2024) for details.

### Volumetric cell segmentation

For normal reproductive stages, we used volumetrically segmented cell reconstructions published by (Witvliet et al. 2021) (L1: Dataset 2, L2: Dataset 5, L3: Dataset 6, Adult: Dataset 8). The segmentation images were acquired from BossDB using *intern* library (Hider et al. 2022; Matelsky et al. 2021). For body wall muscle segmentation, an annotator painted body wall muscles in every 8th section. Then, the intermediate sections were filled by the painted result of the closest section.

Volumetric cell segmentation for dauer images were manually segmented by painting individual sections using VAST (Berger, Seung, and Lichtman 2018). Unlike normal reproductive stages, body wall muscles were painted in every section like other cells in the volume.

### Synapses and connectivity graph

For normal reproductive stages, we used synapses and connectivity information published by Witvliet et al. (Witvliet et al. 2021), publicly available at BossDB (Hider et al. 2022).

### Ground truth annotation for mitochondria detection

For normal reproductive stages and the dauer stage EM image stacks, we picked 10 to 20 sections from each stage. All mitochondria in each section were manually labeled using VAST (Berger, Seung, and Lichtman 2018) and saved as binary images. For dauer stage, synapses and connectivity information generated in Yim et al. (Yim et al. 2024) were used.

### Cell type classification

Cell types based on neuronal function (sensory, inter-, motor neurons) were defined according to classification defined in WormAtlas. When there are multiple types per neuron, we defined it as its primary role. The cell type classifications we have used are consistent with what we have used in Yim et al. (Yim et al. 2024).

Cell types based on the neurotransmitter types (acetylcholine, glutamate, dopamine, GABA, octopamine) were defined according to classification defined in Hobert et al. (Hobert, Glenwinkel, and White 2016). Only primary neurotransmitters were used for classification.

### Mitochondria reconstruction

In every EM image section, mitochondria were detected using deep learning. The network used for the mitochondria detection was adopted from 2D symmetric U-Net architecture (Ronneberger, Fischer, and Brox 2015). The architecture was composed of five layers with a number of feature maps 16, 32, 64, 128, 256 from the top most layer to the bottom most layer, respectively. For each step in downsampling and upsampling layers, the architecture consisted of three 3 x 3 non-strided same convolutions. In the downsampling layers, max pooling was used to downsample by a factor of 2 in each layer. In the upsampling layers, a transposed convolution layer with nearest neighbor interpolation followed by 2 x 2 non-strided convolution was used to upsample by a factor of 2 in each layer. Skip connection was included in every layer which concatenates feature maps at the same level of the left hand side to the output of the transposed convolution layer. Instance normalization (Ulyanov, Vedaldi, and Lempitsky 2016) and rectified linear unit (ReLU) was added after each 3 x 3 convolution. At the end of the symmetric network, 3 x 3 non-strided same convolution and sigmoid function was applied to produce the same-sized output image, where each pixel value represents probability of each pixel belonging to mitochondria.

From the probability map, we thresholded the image with a pixel threshold of 128 / 255 (50%) and generated binarized prediction images of mitochondria. To reconstruct 3D mitochondria, we applied connected components with connectivity of 26. The errors in resulting reconstructions were corrected manually. All the mitochondria were skeletonized using Kimimaro (Silversmith et al. 2022, 2021).

### Computation of synaptic properties

Out– and in-degrees were calculated by counting the total number of postsynaptic and presynaptic partners respectively. For example, if there’s 3 postsynaptic partners for an active zone, 3 will be added to compute out-degree. Synapse size is approximated by measuring the size of the active zone. We count the number of voxels, then multiply by the voxel resolution to get the physical size of an active zone. In this paper, the terms “active zone size” and “synapse size” are used interchangeably. Fan-out is a measure that is computed for each active zone, indicating the number of postsynaptic partners per active zone. Cells with either 0 out-or in-degree are neglected from the corresponding analyses.

### Computation of mitochondria properties

Number of mitochondria is the total number of mitochondria in a cell. Mitochondria volume indicates the size of each mitochondria. The volume is calculated by multiplying the voxel count by the voxel resolution. For analyses done per cell, mitochondria volume means total sum of volumes of all mitochondria included in a cell (Figures 6-8). Mitochondria volume fraction is a proxy for mitochondria density of each cell. The volume fraction is computed by dividing the total mitochondria volume in a cell by the volume of a cell. Mitochondria surface area per volume (SAV) is calculated by dividing the total sum of surface area of all mitochondria by the total sum of volumes of all mitochondria. Mitochondrial complexity index (MCI; Vincent et al. 2019) was computed as

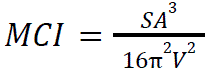

where *SA* is the surface area and *V* is the volume of mitochondria. Cells without any mitochondria have been neglected from the corresponding analyses.

### Computation of distance between synapses and mitochondria

Mitochondria positions were determined by the centroids of segmented mitochondria and the positions of presynaptic sites were determined by the centroids of segmented active zones. For postsynaptic sites, the positions were assigned to the active zone positions for corresponding presynaptic partners. Depending on the analysis, we either measured distance to nearest mitochondria from pre-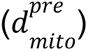 and postsynaptic 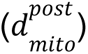 sites, or distance from each mitochondria to nearest pre-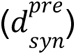 and postsynaptic 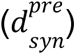 sites by calculating the euclidean distance between two positions.

For randomizations used in Figures 2C and 2D, we assigned random mitochondria within the same neuron instead of the nearest mitochondria for each active zone and measured the distance. 1000 random configurations have been generated and mean and 95% confidence intervals are shown in Figure 2C. In Figure 2D, distribution of distances from one randomly selected configuration is shown as representative.

To distinguish nearby and far mitochondria from synapses as in Figures 3A and 3B, we set the threshold to be the distance where the cumulative synapse distribution starts to saturate (<5% difference). The resulting threshold distances were 1.3 µm (L1), 1.3 µm (L2), 1.1 µm (L3), 1.1 µm (adult), and 1.1 µm (dauer).

### Microdroplet swimming assay

A microdroplet swimming assay was performed to simultaneously measure the frequency of reorientation in a large number of nematodes. Young adult animals were grown on standard nematode growth medium plates containing E. coli OP50. Nematodes in the food lawn were washed in M9 solution to remove *E*. *coli* attached to their bodies, and their behavior was measured by placing them in 1.5 microliters of M9. Swimming and reorientation behavior videos were recorded at 10 Hz. Custom algorithm was used to measure turning rates in micro-droplets (Choi et al. 2020). The experiments were only conducted in the adult stage since turns are difficult to detect in other developmental stages as the worms are small. Besides, the worms tend to stay motionless in the dauer stage.

### Principal component analysis of mitochondrial features

To investigate the overview of mitochondria features in neurons, 8 structural features of mitochondria were used: Total mitochondria volume, mean mitochondria volume, total mitochondria surface area, mean mitochondria surface area, total mitochondria length, mean mitochondria length, mitochondria volume fraction, and mean mitochondrial complexity index. The features were standardized and projected onto the first two principal components.

### UMAP analysis of mitochondrial features

To investigate the overview of mitochondria features in body wall muscles, 5 structural features of mitochondria were used: Total mitochondria volume, total mitochondria surface area, mitochondria volume fraction, mitochondria surface area per volume, and mitochondrial complexity index. The features were standardized and projected onto the first two components of UMAP embeddings (McInnes, Healy, and Melville 2018).

### Sample preparation and fluorescence imaging

Confocal microscopy (ZEISS LSM700; Carl Zeiss) was used to observe mitochondrial transgene expression in the body wall muscle of *C. elegans*. For microscopy and imaging, transgenic animals were paralyzed with 3 mM levamisole and mounted on 3% agar pads. All transgenic animals were observed and imaged at the stage as described: L1, L2, L3, L4, Day 0 adult, Day 5 adult, dauer, Day 0 postdauer adult, Day 5 postdauer adult. All stage worms were collected after *C. elegans* synchronization (L1) with cultivation during proper developmental time except dauer and postdauer adult.

### Neurons and mitochondria renderings

All the renderings of neurons and mitochondria were created in Python using MeshParty (github.com/CAVEconnectome/MeshParty). Screenshots of Neuroglancer (https://github.com/google/neuroglancer) were used for some figures (Figure S5).

### Statistical testing

To find the significance in linear correlation, Pearson correlation was used (Figures 2 and S2). For comparing two independent distributions, Wilcoxon rank-sum test was used as most of the distributions did not follow the normal distribution (Figures 2-8, S2, S7, S8). Significance tests with *p* values less than 10^−20^ were marked as *p*≈0.

### Data and code availability

Mitochondria reconstruction and analysis code is available at https://github.com/jabae/Cmito. EM images, cell segmentation images, and connectivity data (Witvliet et al. 2021; Yim et al. 2024) are available in BossDB. Mitochondria reconstructions of all development stages are available upon request.

