## Supplemental Figures for "Structural diversity of mitochondria in the neuromuscular system across development"

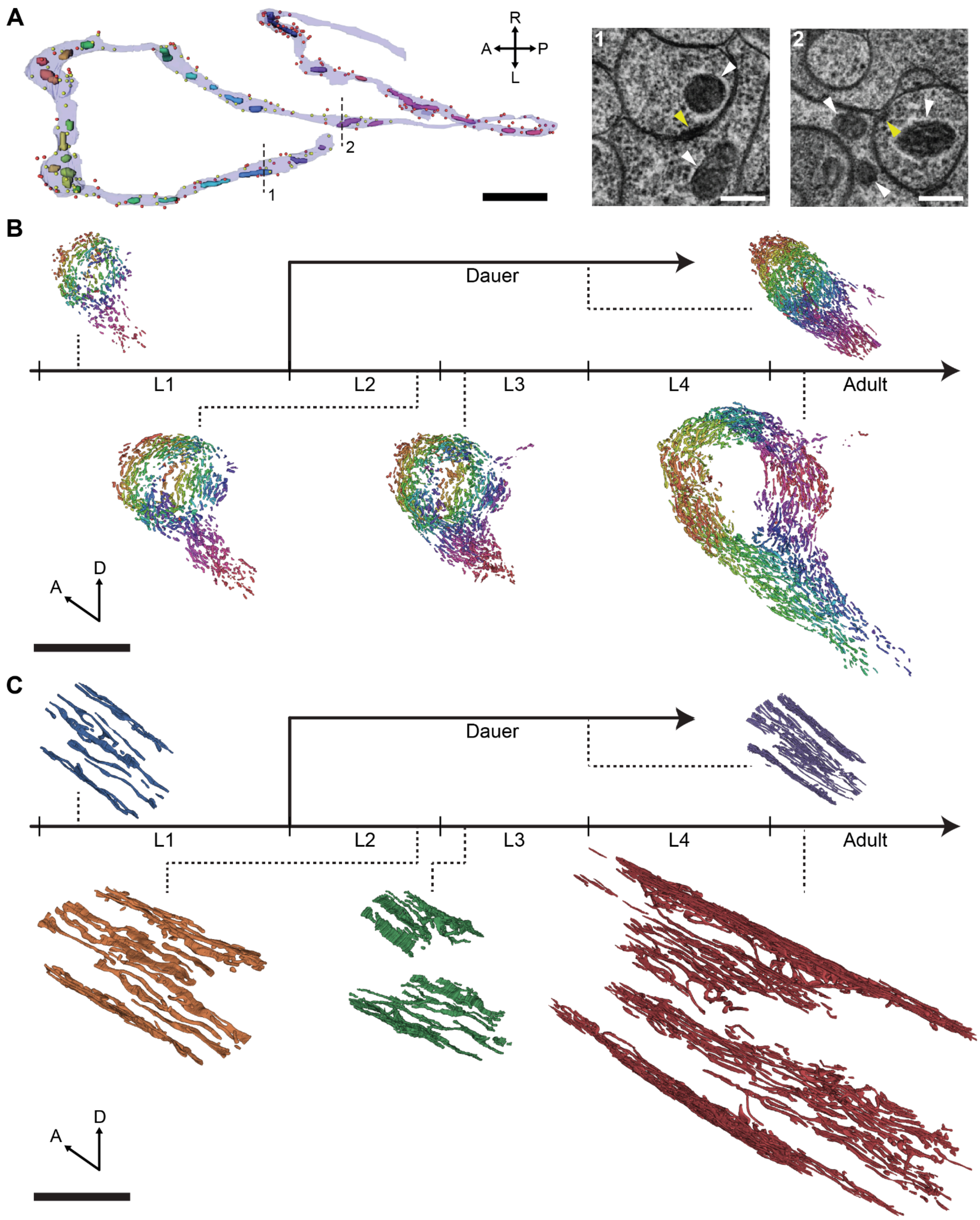


**Figure S1. Mitochondria reconstructions, related to Figure 1.** (A) Mitochondria reconstruction pipeline. Pixels included in the mitochondria are predicted from EM images using convolutional neural network. Then, distinct mitochondria are separately segmented and each segment is meshed for 3D visualization. (B) Total number of neuronal mitochondria in the nerve ring across development. (C)

Individual neuronal mitochondrion volume across development. (D) Individual neuronal mitochondrion length across development. (E) Sum of all neuronal mitochondria volume in the nerve ring across development. (F) Individual neuron volume across development. (G) Sum of all neuron volume across development.
(A) Scale bar: 200 nm. (C, D, F) Mean±stderr.

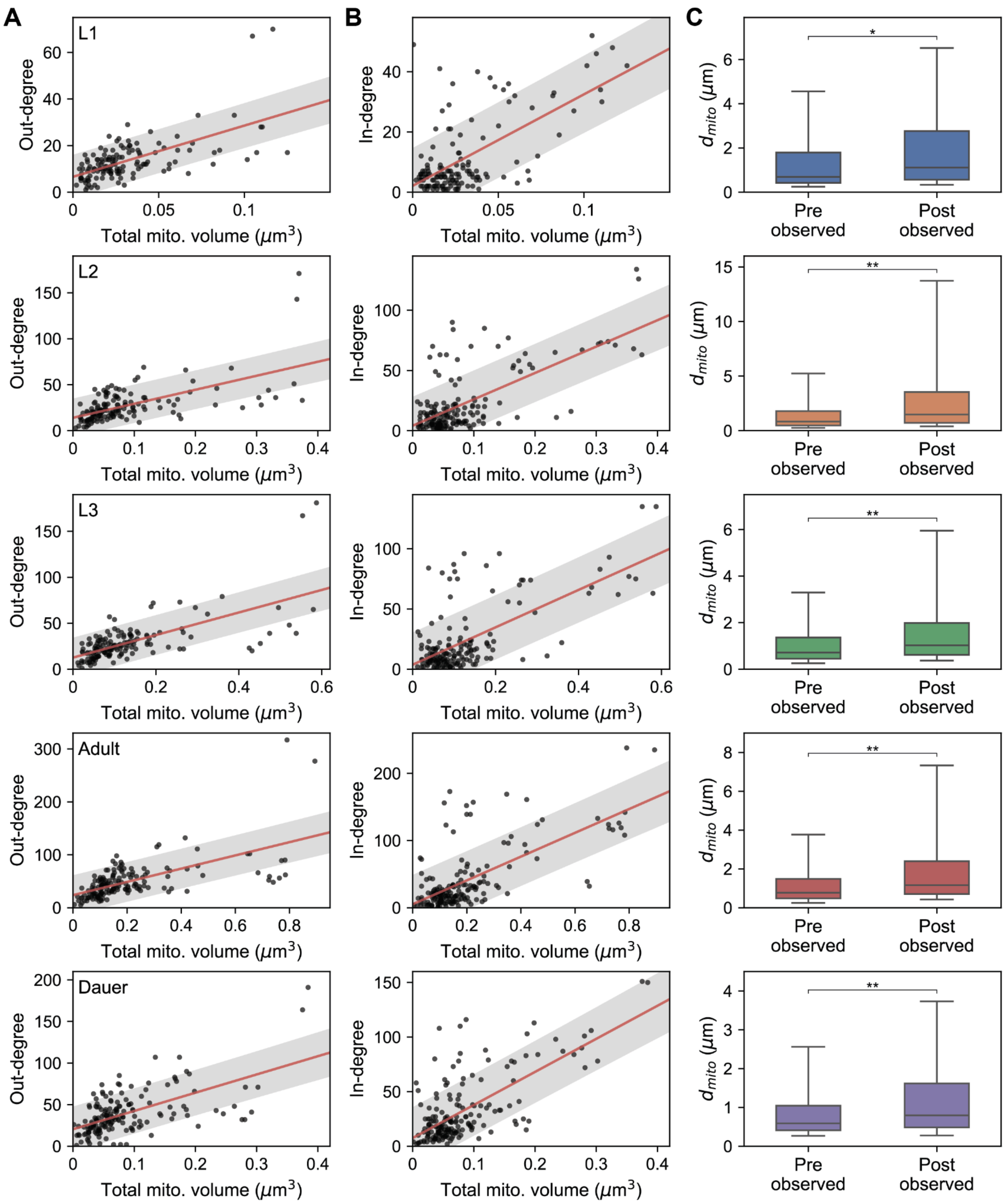


**Figure S2. Mitochondria are spatially correlated with synapses, related to Figure 2.** (A) Relation between mitochondria volume and out-degree across development (L1: n=135, $r$=0.62, $p$=6.96x10$^{-16}$; L2: n=150, $r$=0.60, $p$=4.25x10$^{-16}$; L3: n=152, $r$=0.66, $p$=1.51x10$^{-20}$; adult: n=161, $r$=0.62, $p$2.46x10$^{-18}$; dauer: n=164, $r$=0.59, $p$=4.68x10$^{-17}$; Pearson correlation). (B) Relation between mitochondria volume and in-degree across development (L1: n=140, $r$=0.64, $p$=1.93x10$^{-17}$; L2: n=168, $r$=0.67, $p$=1.93x10$^{-23}$; L3: n=165, $r$=0.66, $p$-=5.42x10$^{-22}$; adult: n=179, $r$=0.68, $p$=2.16x10$^{-25}$; dauer: n=178, $r$=0.69, $p$=1.22x10$^{-26}$;

Pearson correlation). (C) Distributions of distance to nearest mitochondria from pre- and postsynaptic sites (L1: $n_{pre}$=796, $n_{post}$=1568, $p=1.44\times10^{-20}$; L2: $n_{pre}$=1879, $n_{post}$=4033, $p\approx0$; L3: $n_{pre}$=2005, $n_{post}$=3844, $p\approx0$; adult: $n_{pre}$=3677, $n_{post}$=7099, $p\approx0$; dauer: $n_{pre}$=2813, $n_{post}$=5707, $p\approx0$).
(C) Center line: median, box: interquartile range, whiskers: 5th and 95th percentile. *$p<10^{-19}$, **$p\approx0$; Wilcoxon rank-sum test.

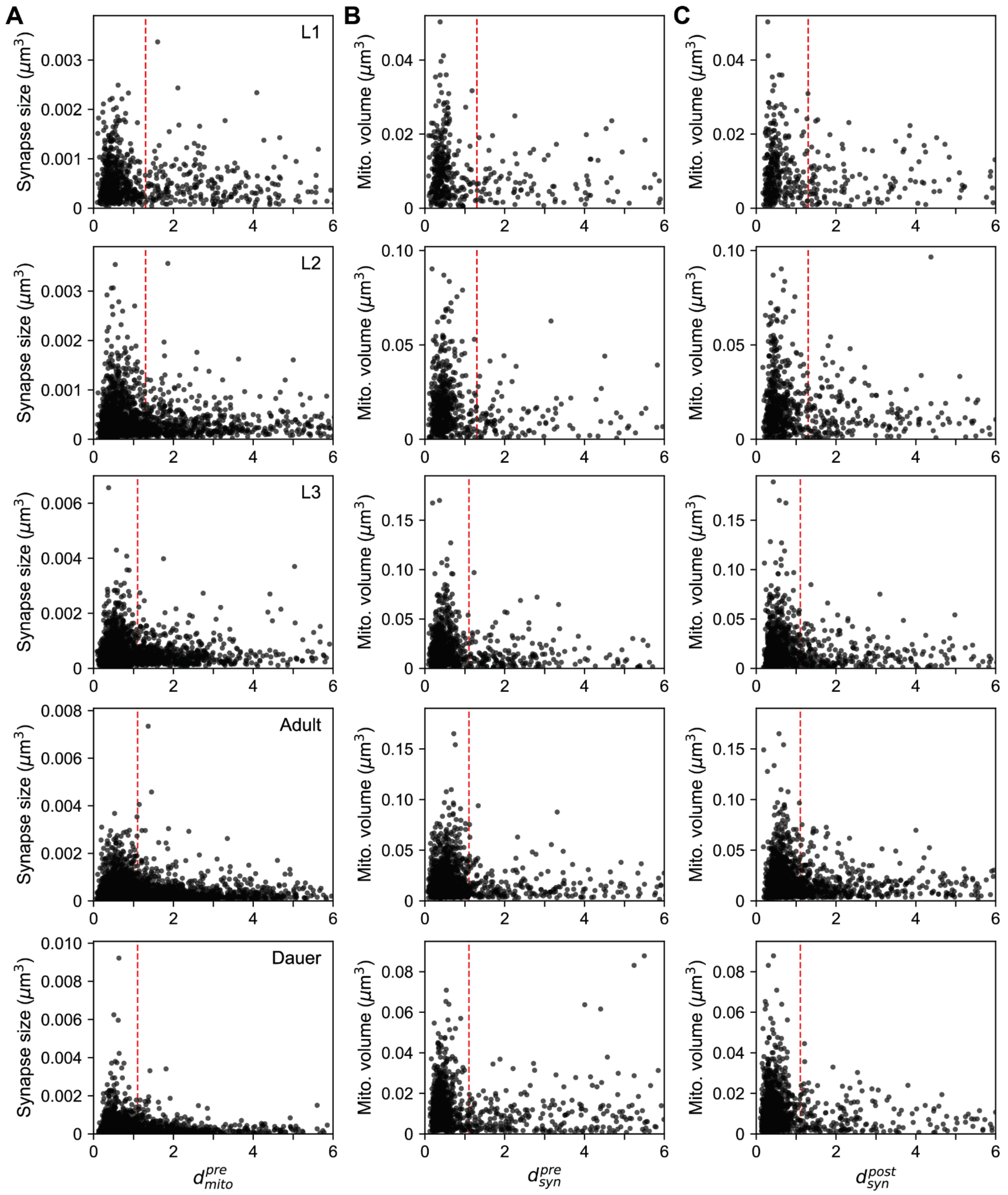


**Figure S3. Larger mitochondria support larger synapses, related to Figure 3.** (A) Relation between distance to nearest mitochondria from each active zone and active zone size across development. (B and C) Relation between distance from nearest mitochondria to pre- (B) and postsynaptic sites (C) and mitochondria volume across development.

(A-C) Red dashed: boundary to classify mitochondria nearby (STAR Methods).

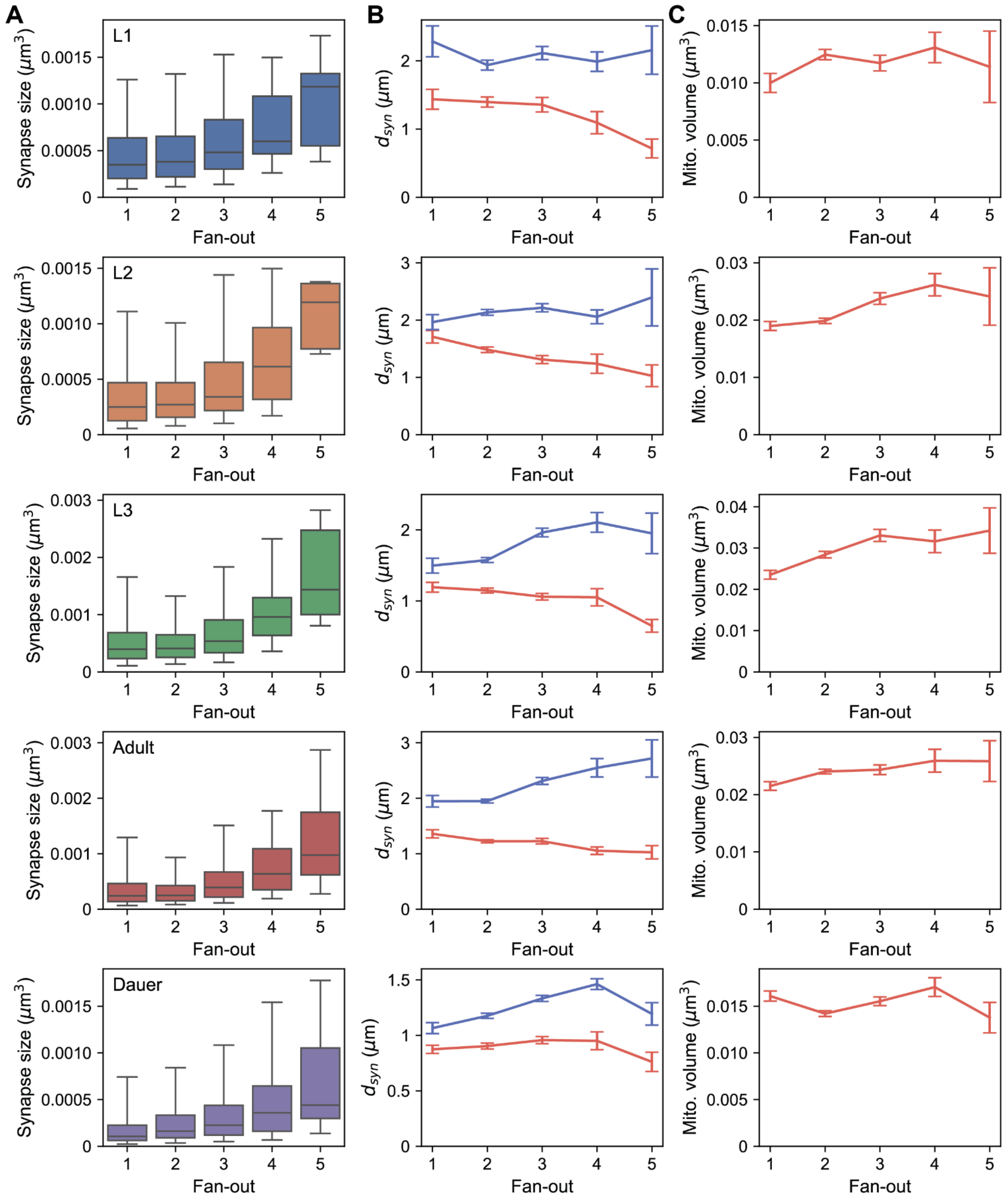


**Figure S4. Larger mitochondria are located closer to support synapses with higher fan-out, related to Figure 3.** (A) Relation between fan-out and active zone size across development. (B) Relation between fan-out and distance from nearest mitochondria to pre- (red) and postsynaptic (blue) sites across development. (C) Relation between fan-out and volume of the nearest mitochondria across development. (A) Center line: median, box: interquartile range, whiskers: 5th and 95th percentile. (B and C) Mean±stderr.

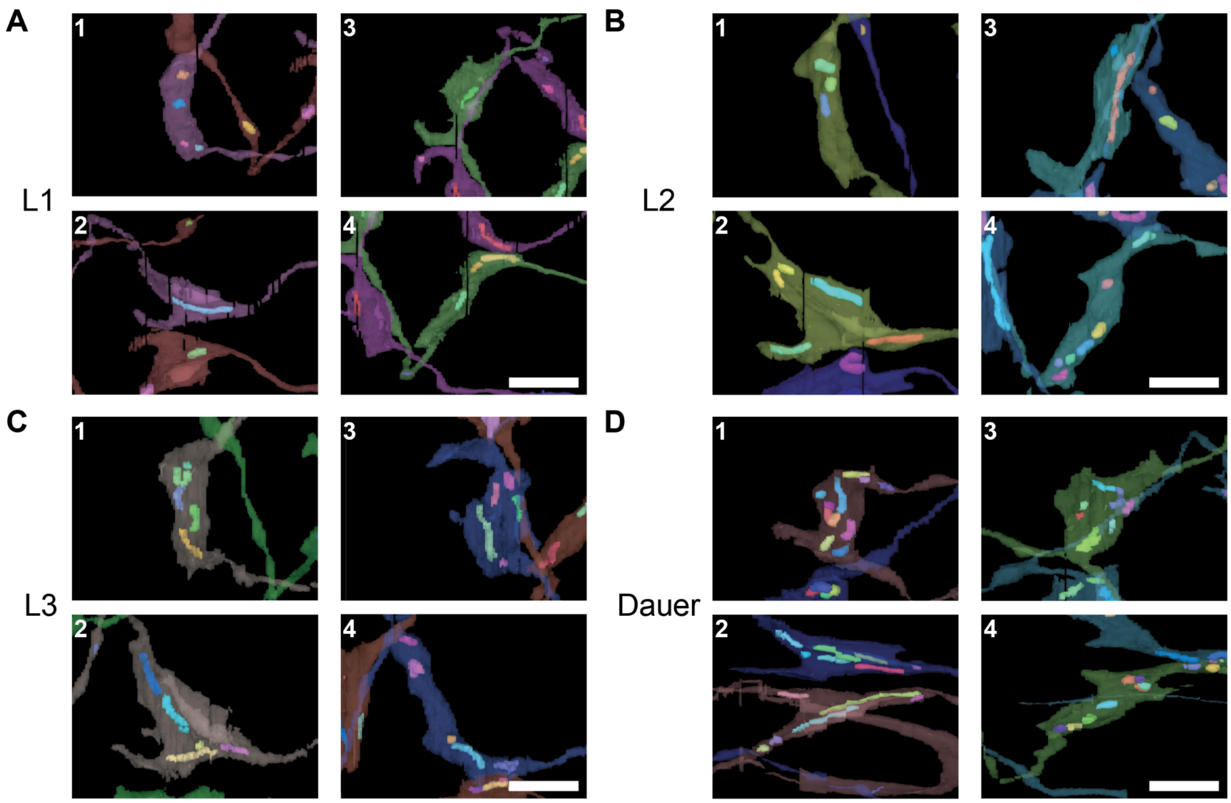


**Figure S5. SMDD and SMDV neurons with mitochondria in other stages, related to Figure 4.** (A-D) Mitochondria in SMDD dorsal (1), SMDD ventral (2), SMDV dorsal (3), and SMDV ventral (4) boutons in L1 (A), L2 (B), L3 (C), and Dauer (D).
(A-D) Scale bars: 2 μm.

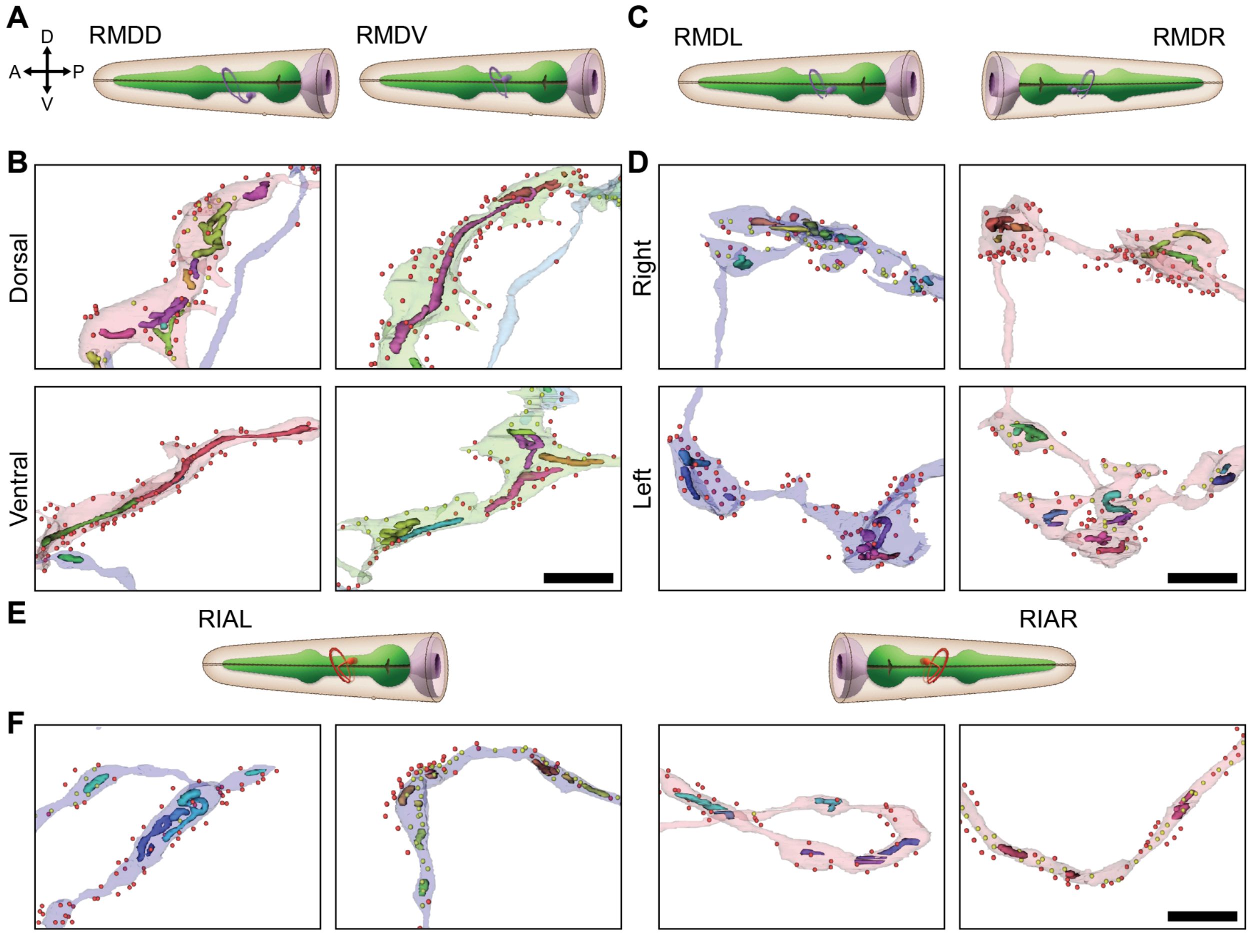


**Figure S6. Neurons with compartmentalized synapse distribution, related to Figure 4.** (A) Diagram of RMDD (left) and RMDV (right) neurons. (B) Reconstructed mitochondria in adult RMDD (left) and RMDV dorsal (top) and ventral (bottom) boutons. Presynaptic (yellow) and postsynaptic (red) sites are marked with dots. (C) Same with (A) for RMDL (left) and RMDR (right) neurons. (D) Reconstructed mitochondria in adult RMDL (left) and RMDR (right) right (top) and left (bottom) boutons. (E) Same with (A) for RIAL (left) and RIAR (right) neurons. (F) Reconstructed mitochondria in adult RIAL (purple) and RIAR (pink) dendritic (left) and axonal (right) boutons.
(B, D, and F) Scale bars: 2 μm.

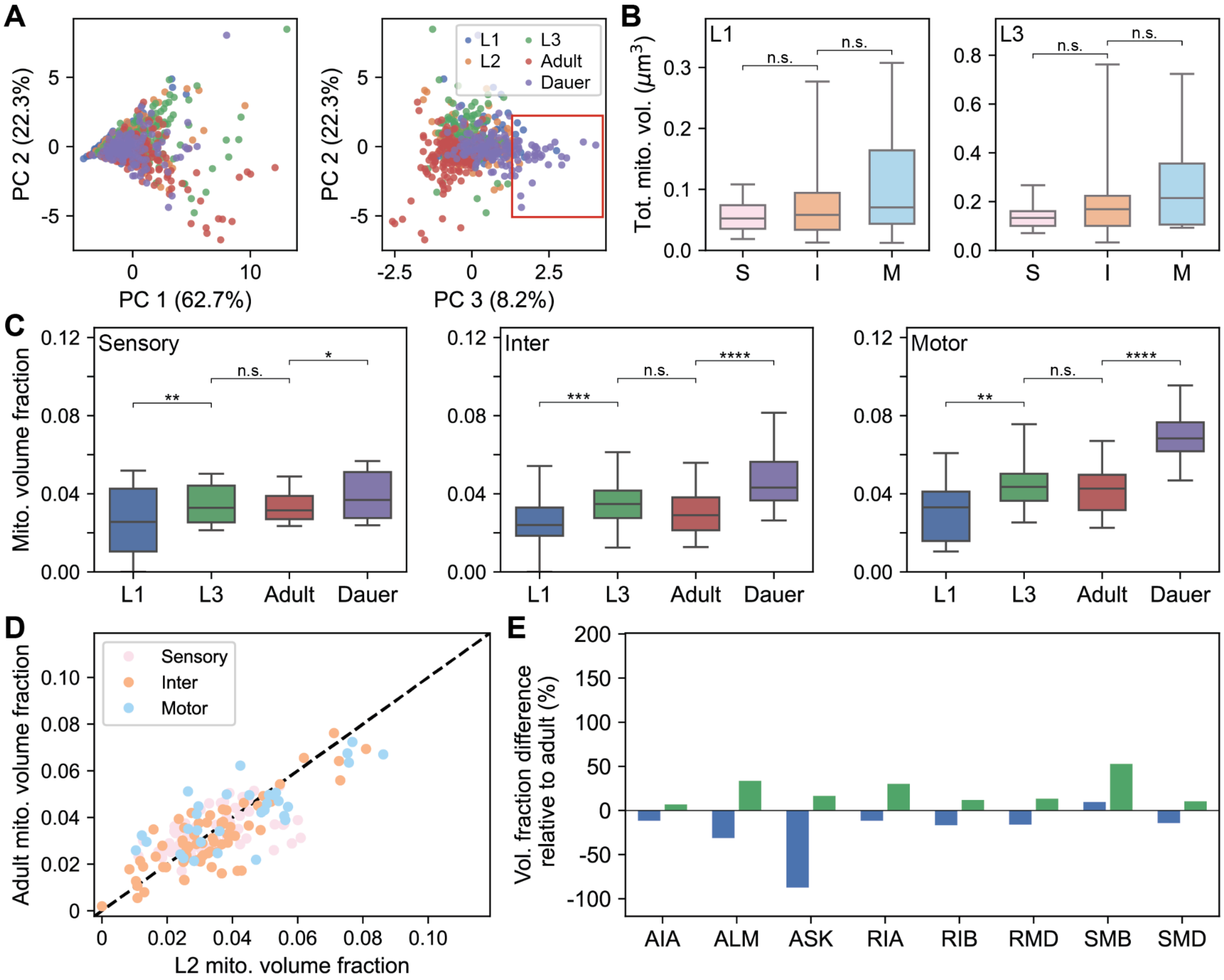


**Figure S7. Neuronal mitochondria features in other stages, related to Figure 6.** (A) Principal component embeddings of neuronal mitochondria features show group of dauer neurons clustered outside (red box). (B) Distribution of total mitochondria volume in neurons per type for L1 (left; $p$=0.35, $p$=0.15) and L3 (right; $p$=0.0.12, $p$=0.09). (C) Distribution of mitochondria volume fraction in L1, L3, adult, and dauer for sensory (left; $p$=0.0098, $p$=0.92860, $p$=0.0450), inter- (middle; $p$=1.78x10$^{-4}$, $p$=0.065, $p$=2.99x10$^{-8}$), and motor (right; $p$=0.002, $p$=0.46, $p$=3.69x10$^{-8}$) neurons. (D) Comparison between mitochondria volume fraction of L2 and adult. (E) Difference in mitochondria volume fraction compared to selected neurons in adult for L1 (blue) and L3 (green).

(B and C) $n_{sen}$=61, $n_{int}$=69, $n_{mot}$=32. Center line: median, box: interquartile range, whiskers: 5th and 95th percentile. *$p$<0.05, **$p$<0.01, ***$p$<0.001, ****$p$<10$^{-7}$; Wilcoxon rank-sum test.

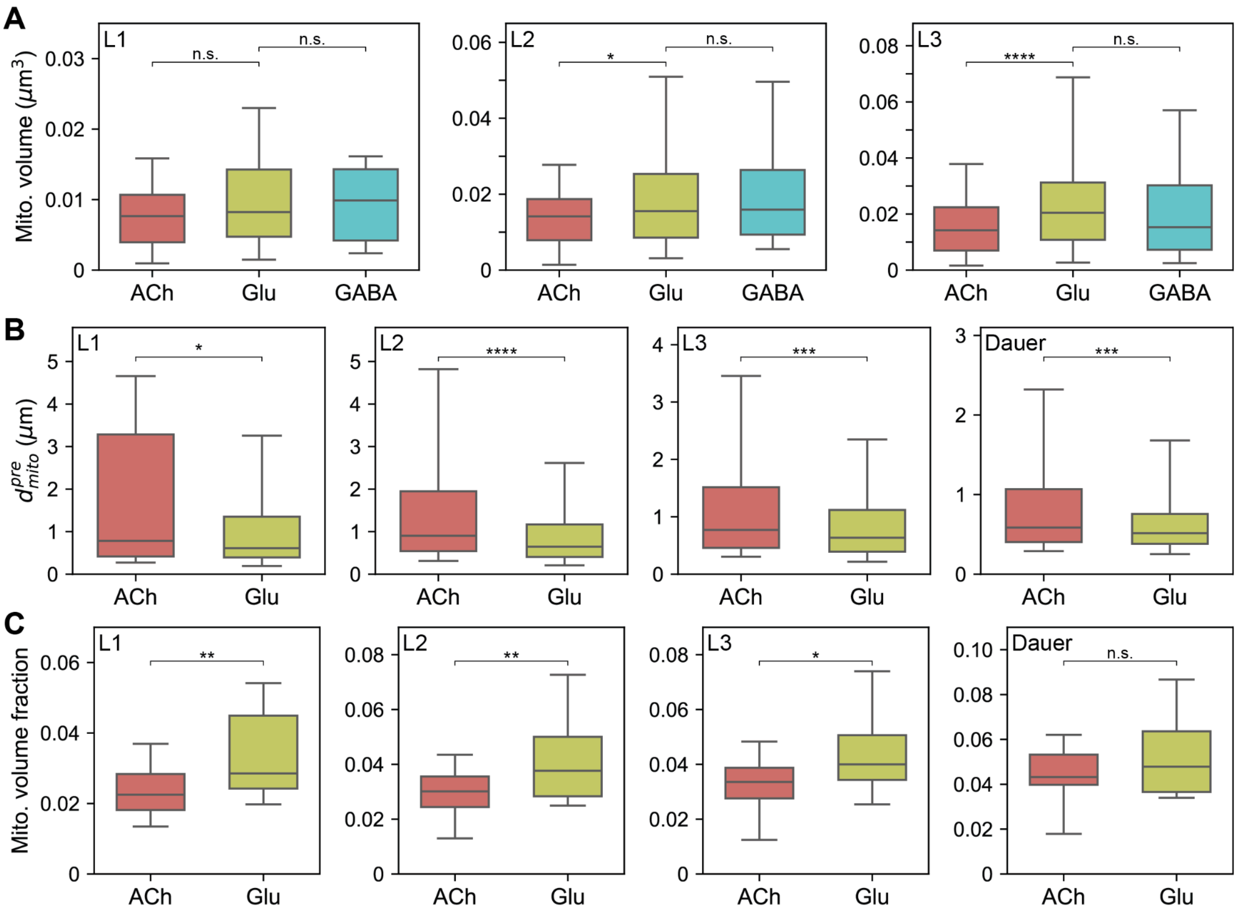


**Figure S8. Mitochondria features for different neurotransmitters in other stages, related to Figure 7.** (A) Distributions of mitochondria volume in interneurons that release acetylcholine (red), glutamate (green), and GABA (blue) in L1 (left; $n_{ACh}$=65, $n_{Glu}$=98, $n_{GABA}$=25, $p$=0.095, $p$=0.890), L2 (middle; $n_{ACh}$=115, $n_{Glu}$=147, $n_{GABA}$=40, $p$=0.024, $p$=0.727), and L3 (right; $n_{ACh}$=161, $n_{Glu}$=172, $n_{GABA}$=57, $p$=7.48x10$^{-5}$, $p$=0.256). (B) Distribution of distance to the nearest mitochondrion from each active zone in interneurons that release acetylcholine (red), glutamate (green) in L1 ($n_{ACh}$=106, $n_{Glu}$=205, $p$=0.011), L2 ($n_{ACh}$=226, $n_{Glu}$=440, $p$=5.07x10$^{-8}$), L3 ($n_{ACh}$=253, $n_{Glu}$=477, $p$=3.62x10$^{-4}$), and dauer (from left; $n_{ACh}$=304, $n_{Glu}$=567, $p$=2.07x10$^{-4}$). (C) Distribution of mitochondria volume fraction of interneurons that release acetylcholine (red), glutamate (green) in L1 ($n_{ACh}$=26, $n_{Glu}$=20, $p$=0.007), L2 ($n_{ACh}$=28, $n_{Glu}$=20, $p$=0.006), L3 ($n_{ACh}$=28, $n_{Glu}$=20, $p$=0.016), and dauer (from left; $n_{ACh}$=30, $n_{Glu}$=20, $p$=0.362).

(A-C) Ach: Acetylcholine, Glu: Glutamate. Center line: median, box: interquartile range, whiskers: 5th and 95th percentile. *$p$<0.05, **$p$<0.01, ***$p$<0.001, ****$p$<10$^{-4}$; Wilcoxon rank-sum test.

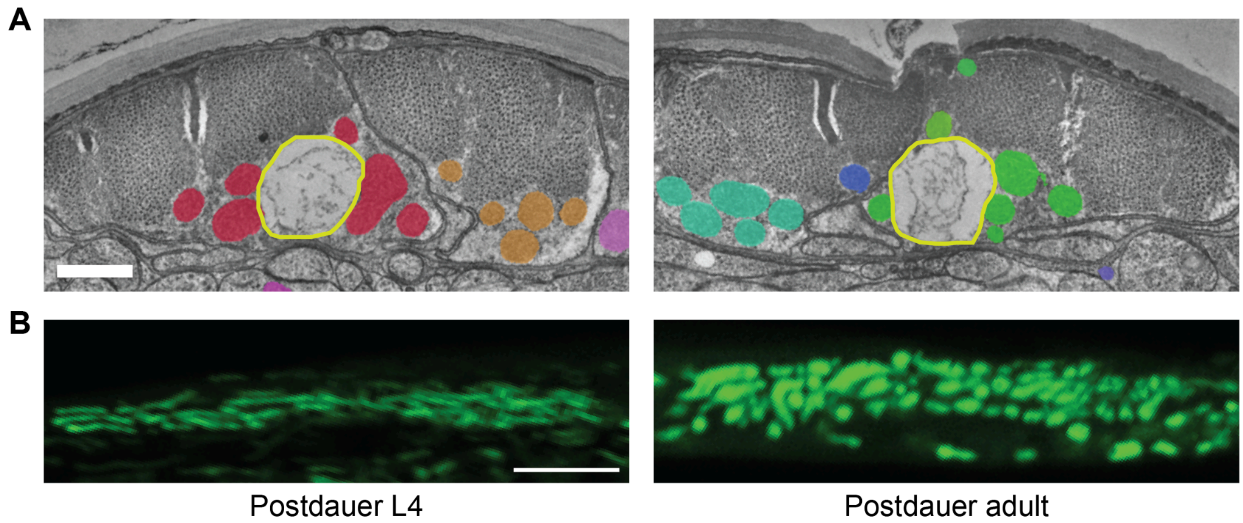


**Figure S9. Mitochondria in the body wall muscles, related to Figure 8.** (A) Lipid droplets (yellow) occupy the space in between the mitochondria strands (color shades) in the body wall muscles of dauer, serving as an energy source. (B) Fluorescence images of BWM mitochondria in postdauer L4 (left) and postdauer adult (right).

(A) Scale bar: 500 nm. (B) Scale bar: 5 µm.
